# Circulating, cell-free methylated DNA indicates cellular sources of allograft injury after liver transplant

**DOI:** 10.1101/2024.04.04.588176

**Authors:** Megan E. McNamara, Sidharth S. Jain, Kesha Oza, Vinona Muralidaran, Amber J. Kiliti, A. Patrick McDeed, Digvijay Patil, Yuki Cui, Marcel O. Schmidt, Anna T. Riegel, Alexander H.K. Kroemer, Anton Wellstein

**Author notes:** **Corresponding author**: Anton Wellstein - Contact; Mailing address: Georgetown University Medical Center, 3970 Reservoir Road, New Research Building E311, Washington, D.C. 20057 USA.

## Abstract

Post-transplant complications reduce allograft and recipient survival. Current approaches for detecting allograft injury non-invasively are limited and do not differentiate between cellular mechanisms. Here, we monitor cellular damages after liver transplants from cell-free DNA (cfDNA) fragments released from dying cells into the circulation. We analyzed 130 blood samples collected from 44 patients at different time points after transplant. Sequence-based methylation of cfDNA fragments were mapped to patterns established to identify cell types in different organs. For liver cell types DNA methylation patterns and multi-omic data integration show distinct enrichment in open chromatin and regulatory regions functionally important for the respective cell types. We find that multi-tissue cellular damages post-transplant recover in patients without allograft injury during the first post-operative week. However, sustained elevation of hepatocyte and biliary epithelial cfDNA beyond the first week indicates early-onset allograft injury. Further, cfDNA composition differentiates amongst causes of allograft injury indicating the potential for non-invasive monitoring and timely intervention.

## Introduction

Liver transplant is the standard-of-care for patients with end-stage liver disease and is the second most common transplant after the kidney (1). Despite improved survival rates, there is still a high prevalence of complications contributing to perioperative mortality post-liver transplant, mostly occurring within the first month (2, 3). Unfortunately, current non-invasive biomarkers have a limited scope and fail to identify cellular causes of allograft injury (4). Thus, tissue biopsy is still the gold-standard to confirm diagnosis and monitor response to treatment. The analysis of cell-free DNA (cfDNA) in the circulation is an alternative to conventional biomarkers. CfDNA consists of fragments shed by dying cells throughout the body and its analysis can serve as a non-invasive approach for monitoring allograft as well as host tissue changes at a cellular level following liver transplant (5–11). Genetic differences between donor and recipient (SNPs) have been used to identify donor-derived cfDNA (dd-cfDNA) originating from the transplanted allograft to serve as a predictive biomarker of allograft injury and rejection (8, 12–18). However, there are many situations when genetic differences cannot be used to identify allograft-derived DNA; for example, when the genotype is unknown, multiple genotypes exist in the host, and when the donor is closely related to the recipient (13). Instead, epigenetic modifications can be used to identify cfDNA that is recipient or allograft-derived by using tissue- and cell-type specific marks that are independent of genotype differences between the donor and recipient (19–29). Allograft injury can thus be detected in an organ specific fashion, which is of critical importance in recipients of multi-organ transplants and recipients of hematopoietic cell transplant (HCT) who develop Graft-versus-Host disease (GvHD) (30–32). Also, the allograft as well as recipient organs are impacted by the transplant process as well as by subsequent treatments that can lead to tissue damage and remodeling. Primary injuries or secondary changes stemming from tissue repair can be quantified from cfDNA to indicate cell-type-specific damage (1–4, 33, 34).

DNA methylation is highly cell-type-specific and has been found to reveal the origins of tissue damages and altered cell turnover from cfDNA samples in a wide-range of applications (27, 28, 30, 31, 35–45). DNA methylation patterns are stable epigenetic marks for cells maintained throughout DNA replication and cell proliferation (46). Cell-type-specific DNA methylation established early-on during development has been found to be highly conserved across individuals, irrespective of age and disease status (21, 22, 46–50). Also, cell-free methylated DNA can reflect intervention-related changes over time through analyses of serially collected blood samples (38). Fragment-level deconvolution of methylation sequencing data allows for increased sensitivity and specificity of signal localization using CpG pattern analysis of individual cfDNA molecules (35, 38, 51–53). Likewise, hybridization capture to CpG-rich DNA segments maximizes sequencing depth while still maintaining comprehensive coverage (38).

Here, we utilize circulating, cell-free methylated DNA to monitor cellular damages after liver transplant, impacting the allograft tissue as well as the recipient’s organs. We expand existing cell-type-specific DNA methylation atlases to include non-parenchymal cell-types from the liver, including hepatic stellate, endothelial, liver-resident immune, and biliary epithelial cells. Then, we perform capture-sequencing of cell-free methylated DNA from serial blood samples and evaluate multi-tissue cellular damages after liver transplant. We find that sustained elevation of hepatocyte and biliary epithelial cfDNA beyond the first post-operative week is indicative of allograft injury. In addition, we show that there are significant changes in cfDNA composition corresponding to different allograft injury patterns at time of tissue-biopsy-proven diagnosis. Thus, cell-free methylated DNA can non-invasively indicate cellular sources of allograft injury in liver transplant patients.

## Results

### Cellular damages after liver transplant indicated by cell-free methylated DNA in the circulation

To monitor cellular damages after liver transplant, we collected serial serum samples from 28 adult liver transplant patients during the peri-transplant time period and profiled cfDNA methylation from samples at predetermined timepoints up to one month after transplant (n = 100 samples). We also collected samples from patients experiencing complications and added phenotype-matched samples from an additional 16 patients at the time of for-cause liver biopsy (FC-bx) used to identify allograft injury (n = 30 samples). Cell-free DNA fragments isolated from these 130 serum samples were bisulfite treated, enriched for sequences of interest by methylome-wide hybridization capture and subjected to sequence analysis (**Fig. 1**). The tissue and cell type origins of cfDNA fragments in the circulation were mapped to an expanded atlas of cell-type-specific DNA methylation to infer tissue damages and differentiate amongst causes of allograft injury (Methods). Demographic information and clinical characteristics of patients enrolled in this study are in **Supplemental Table 1.**

**Figure 1.**
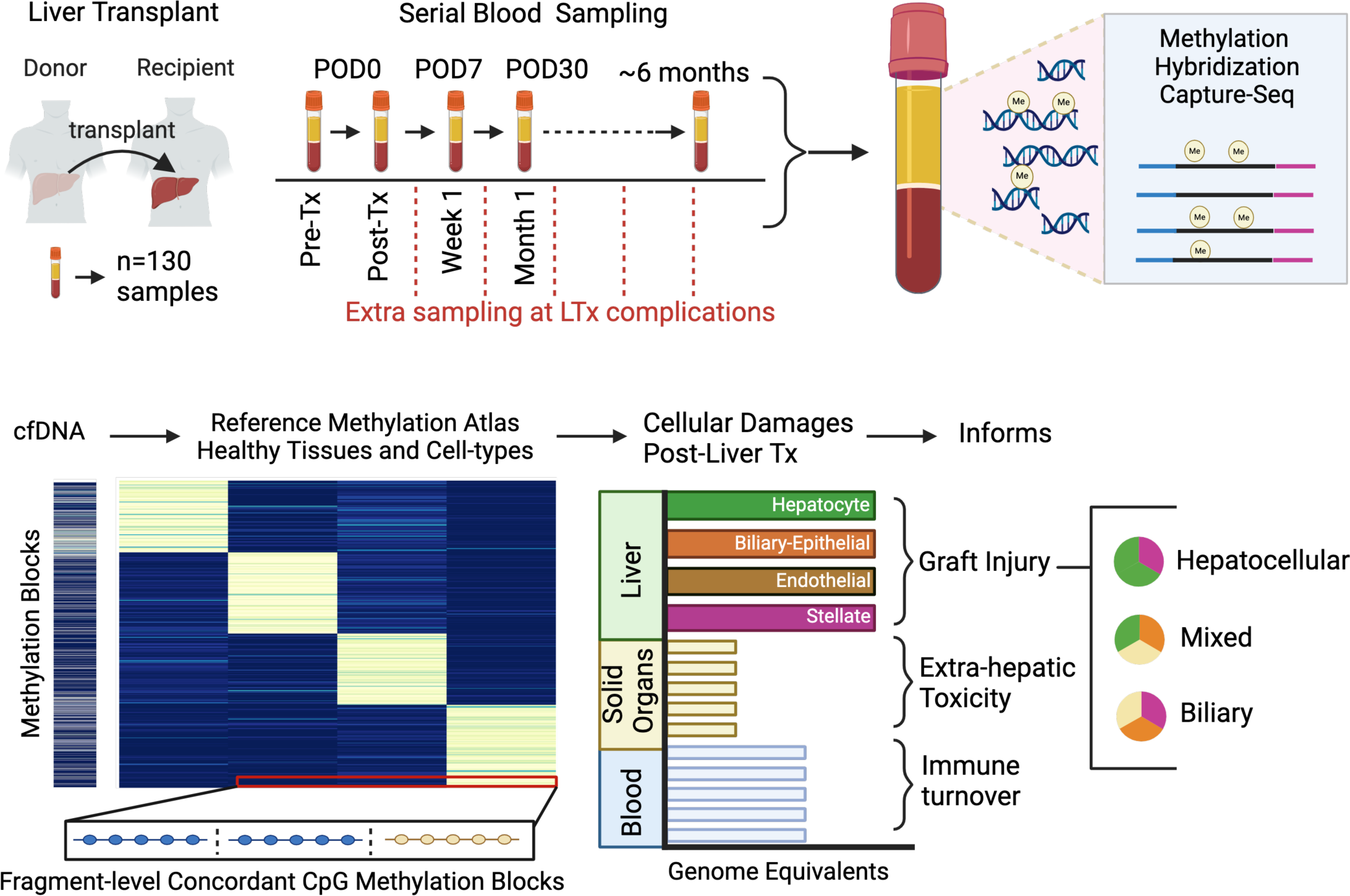
Study overview using cell-free methylated DNA in blood to monitor cellular damages after liver transplant. Serial serum samples were collected from 28 patients before and after liver transplant at predetermined time points (n=100 samples) during the first month with additional timepoints in patients experiencing complications as they arose. We also collected phenotype-matched samples from an additional 16 patients with allograft injury at the time of liver-biopsy proven diagnosis (n=30 samples). Cell-free DNA (cfDNA) methylome profiling of serum samples was performed using hybridization capture-sequencing of bisulfite-treated cfDNA. Then, cell-type-specific DNA methylation blocks identified from reference data of healthy tissues were used to trace the origins of patient cfDNA fragments. Cellular damages of the transplanted organ as well as other recipient organs were quantified to monitor systemic impact.

### Characterization of liver cell-type-specific epigenomes to expand sequencing-based DNA methylation atlas of healthy tissues

To identify cellular origins of cfDNA fragments in the circulation, we expanded the existing cell-type-specific DNA methylation atlas to liver cell-types relevant for injury and repair and generated methylome-sequencing data for hepatic stellate, liver endothelial, biliary epithelial, and liver-resident immune cell populations. In addition, we also included published whole genome bisulfite sequencing (WGBS) data from purified healthy human cell-types (35, 38). This resulted in curation of over 450 WGBS datasets encompassing over 40 cell-types from diverse populations of donors (**Supplemental Table 2**) generating a reference methylome atlas as previously described (35, 38). Briefly, we first segmented the data into homogenously methylated blocks where DNA methylation levels at adjacent CpG sites were highly correlated across different cell types. Then, we restricted the analysis to the 364,268 blocks covered by our hybridization capture panel used in the analysis of cfDNA in human serum (captures 80Mb, ∼20% of CpGs). Average methylation was calculated within blocks of at least three CpG sites and unsupervised clustering analysis was performed for the top 10% variable blocks across all samples. We found that with the additional data incorporated, samples still clustered strongly by cell type and developmental lineage (**Supplemental Fig. 1d**). Notably, parenchymal and non-parenchymal liver cell methylomes did not cluster together. Instead, samples clustered with other cell-types of the same lineage, independent of the germ layer origin of their tissues of residence. Interestingly, biliary epithelial samples isolated from intrahepatic ducts and the gallbladder (columnar epithelium) demonstrated distinct methylation patterns compared to biliary epithelial samples isolated from the larger main hepatic, common bile and pancreatic ducts (cuboidal epithelium) (**Supplemental Figs. 1b and 1c**).

Based on the unsupervised clustering analysis, we grouped the reference WGBS data into 20 groups for downstream analysis (**Supplemental Table 2**). We identified cell-type specific differentially methylated blocks (DMBs) within these groups taking a one-vs-all approach as previously described (38). The co-methylation status of neighboring CpG sites in each block distinguished amongst all cell types included in the final groups (**Supplemental Table 3**). The heatmap in **Fig. 2a** depicts the top 50 blocks with the highest score for each cell-type and the top hepatocyte-specific blocks are emphasized in **Figs. 2b and 2c**. The methylation sequencing approach taken here allowed for assessment of fragment-level methylation patterns rather than the limited single-site resolution of methylation arrays (**Fig. 2d)** (6). Whereas bulk tissue analyses average the methylation status amongst all cell-types, purified cell-specific methylome analysis allowed for discovery of features critical to the identity of non-parenchymal cell-types that contribute only few cells to the overall population and therefore would otherwise be missed (54, 55).

**Figure 2.**
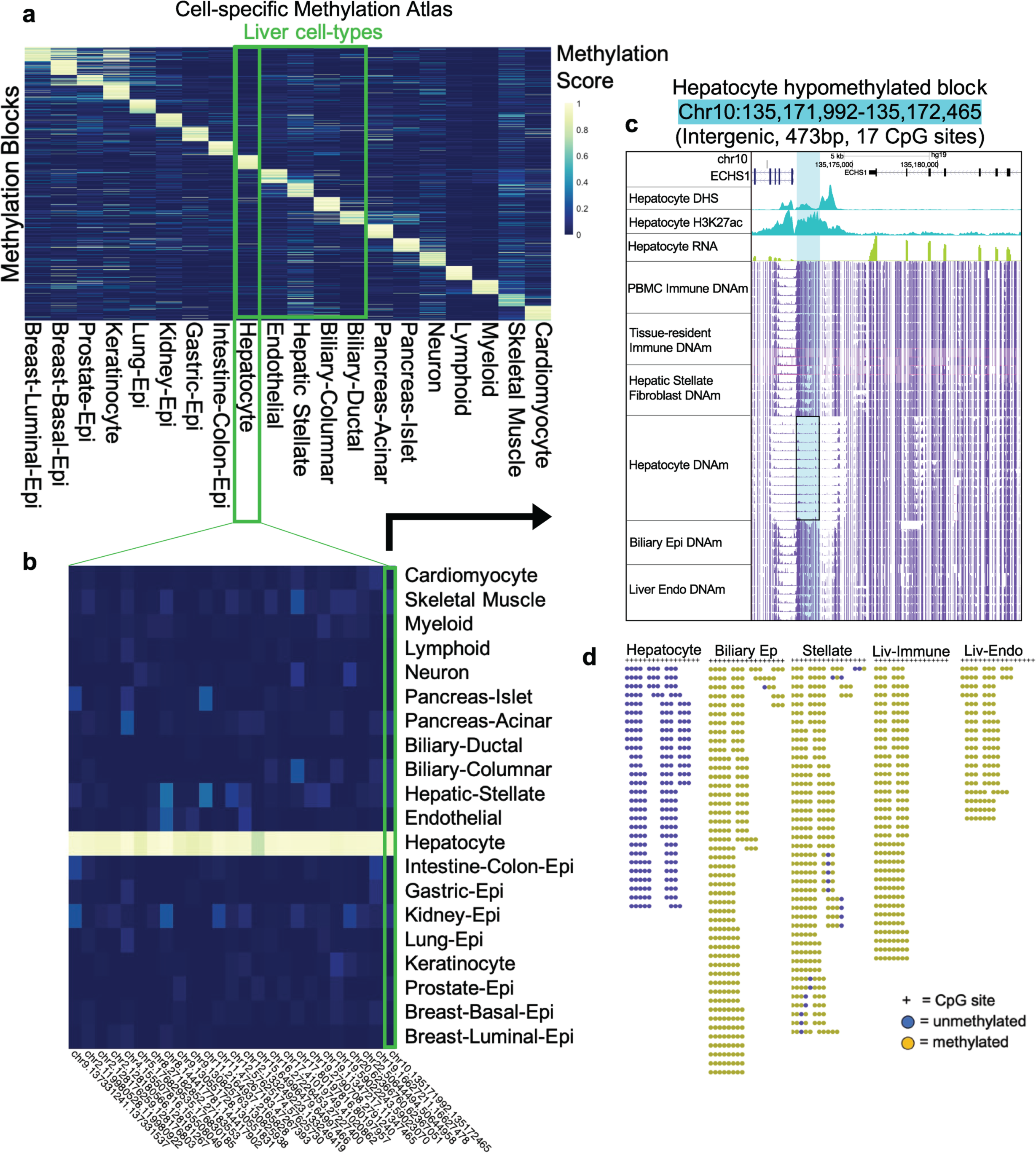
Liver cell-type DNA methylation atlas relative to other healthy tissues. **a**, Heatmap of differentially methylated, cell-type-specific blocks (DMBs) identified from reference WGBS data of healthy human cell-types. Each cell in the plot marks the methylation score of one genomic region (rows) at each of 20 cell-types (columns), with up to 50 blocks shown per cell type. The methylation score represents the number of fully unmethylated or methylated read-pairs divided by total coverage for hypo- and hyper-methylated blocks, respectively. **b**, Heatmap highlighting the top 25 hepatocyte-specific DMBs. **c**, Example of one hepatocyte-specific hypomethylated block (highlighted in blue), upstream of *ECHS1* highly expressed in hepatocytes (green track). The alignment from the UCSC genome browser depicts the average DNA methylation (DNAm, purple tracks) across WGBS samples from five different liver cell-types as well as PBMC samples. Chromatin organization marks in hepatocytes are displayed (blue tracks) to show accessibility (DNAse I hypersensitivity, DHS) and regulatory function (H3K27ac binding). **d**, Fragment-level visualization of methylation sequencing reads at hepatocyte-specific hypomethylated block in reference WGBS samples from five different liver cell-types.

### Liver cell-type-specific hypomethylated blocks coincide with cell-type specific chromatin accessibility and H3K27ac binding

The cell-type-specific DMBs identified using the expanded WGBS reference data resembled those of previously published methylation atlases, being largely hypomethylated, intragenic and annotated to genes relevant for cell function and identity (**Supplemental Figs. 1a and 2f; Supplemental Table 4**) (35, 38, 41). Notably, cell-type-specific hypermethylated DMBs were much less frequent (17% on average) and enriched for CpG islands compared to cell-type-specific hypomethylated DMBs that were located in relatively CpG-depleted, GC-low regions characteristic of programmed demethylation occurring at enhancers (**Supplemental Fig. 2a**) (35, 47, 56). Indeed, the majority of liver cell-type-specific hypomethylated DMBs were enhancers by chromHMM annotations (**Fig. 3c**). In contrast, the majority of liver cell-type-specific hypermethylated DMBs were annotated to bivalent TSS/enhancers and repressive Polycomb targets (**Supplemental Fig. 2d**). This matches with the function of cell-type-specific hypermethylation in repressing genes associated with embryonic stem cell pluripotency to stabilize cellular differentiation during development (**Supplemental Fig. 2e**) (57–59). To further explore the liver cell-type-specific DMBs identified, we generated and compiled additional chromatin accessibility and histone modification data to characterize the integrated epigenomes of hepatocyte, biliary epithelial, hepatic stellate, liver endothelial and liver-resident immune cell-types. We were surprised to find that liver cell-type-specific hypomethylated blocks were also regions with cell-type specific chromatin accessibility and H3K27ac binding, emphasizing the regulatory importance of these regions in maintaining cell-type-specific features reflected in the multi-omic datasets (**Figs. 3a and 3b**).

**Figure 3.**
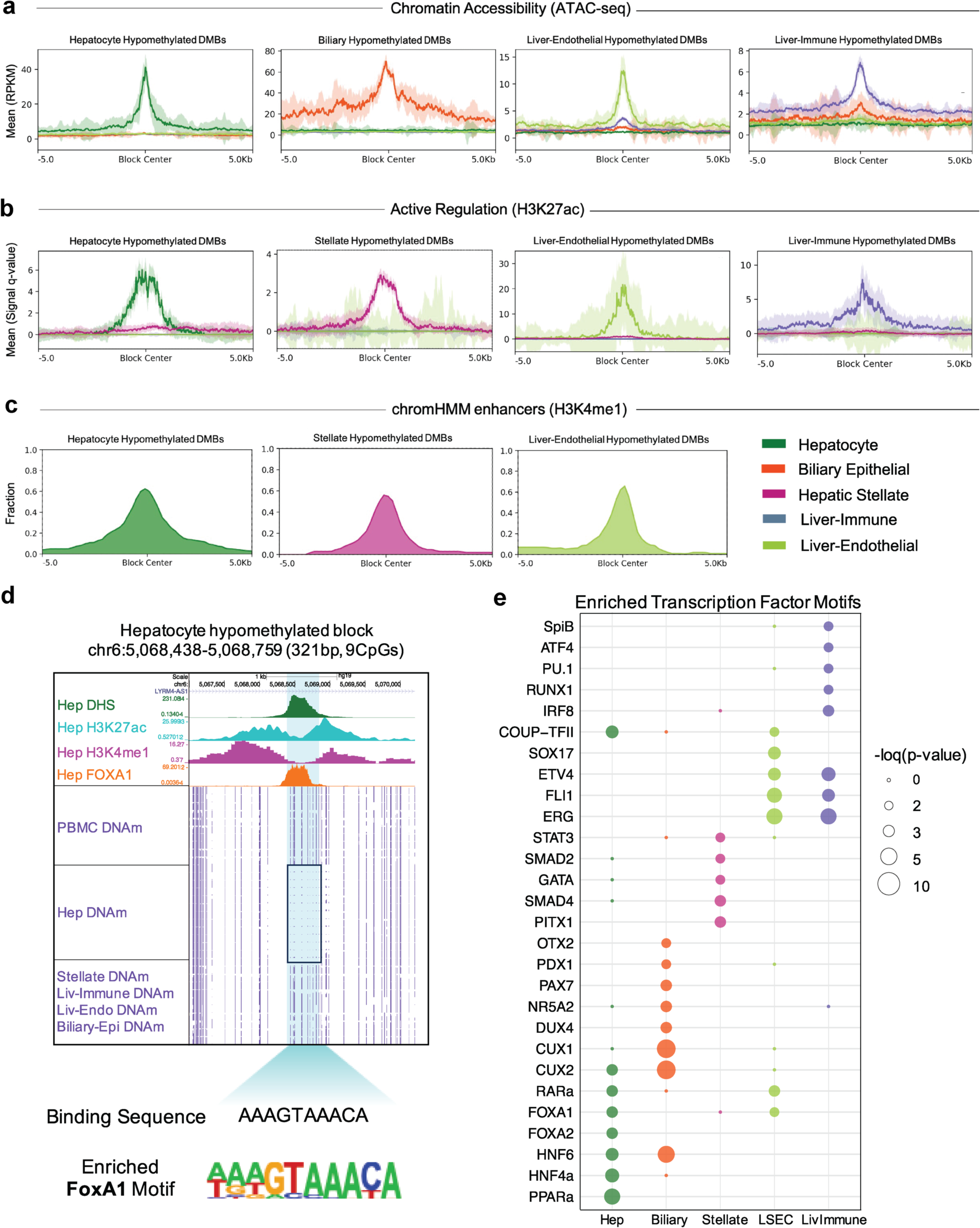
Liver cell-specific hypomethylated DNA blocks coincide with other cell-specific epigenetic marks. **a-b**, Relationship of liver cell-type-specific hypomethylated blocks to chromatin accessibility and H3K27ac binding. Summary plots show the intensity of DHS/ATAC-seq or H3K27ac marks in a ±5-kb regions surrounding each hepatocyte-, biliary columnar epithelial-, hepatic stellate-, liver endothelial-, and liver-immune cell-specific hypomethylated block, respectively. Solid lines represent plot summary with standard error depicted by semi-transparent colored region. **c**, Fraction of cell-type specific hypomethylated blocks labeled as enhancers, associated with H3K4me1 mark, in chromHMM annotations for the same cell-type. **d**, UCSC genome browser alignment at one example hepatocyte-specific hypomethylated block containing the FOXA1 binding sequence. Average methylation across WGBS samples shown in purple tracks. **e**, Pioneer and developmental TF binding sites enriched within liver cell-type specific hypomethylated blocks, from HOMER motif analysis. Captured blocks without liver cell-type-specific methylation were used as background. Hepatocyte DHS, H3K27ac, and H3K4me1 data were obtained from the German Epigenome programme (DEEP). Hepatocyte FOXA1 Chip-seq data was obtained from the ENCODE project. Biliary epithelial ATAC-seq data and Hepatic stellate H3K27ac histone modification data were obtained from the ENCODE project.

### Pioneer transcription factor binding sites (TFBS) enriched within liver cell-type-specific DNA methylation blocks

We performed motif analysis to explore association of the identified liver cell-type-specific DNA methylation with transcription factor binding. We found enriched motifs for several pioneer transcription factors within hypomethylated DMBs, including FOXA1/2, PAX7, CUX1, HNR5A2, DUX4, OTX2, GATA, SOX17, ATF4 and PU.1 (**Figs. 3d and 3e**). We also found enriched motifs of binding sites for several liver developmental TFs known to cooperate with pioneer TFs, including HNF4a, HNF6, PDX1, RARa, COUP-TFII, and RUNX1 (60–62). Pioneer factors are a subclass of TFs that can bind to closed chromatin and elicit an extended functional capacity of the domain, often through local chromatin opening and demethylation (60). As such, they act as master regulators of development and are known to drive cell fate transitions (60). However, we were surprised to find binding sites for pioneer TFs also enriched within liver-specific hypermethylated DMBs (**Supplemental Fig. 2c**). Interestingly, CpG dinucleotides were also enriched within the motifs found in hypermethylated DMBs and several methylation-sensitive TFs were amongst the top hits, including NRF1 where methylation is known to directly repress TF binding (**Supplemental Fig. 2b and Supplemental Table 5**) (63). Although less common, pioneer TFs have been shown to recruit transcriptional repressors and establish a closed and further silenced chromatin architecture (57, 64, 65). Several TFBS of transcriptional repressors known to interact with pioneer TFs were also enriched, including TBET, TRPS1, ZNF669, and E2F7. Annotation of the majority of liver-specific hypermethylated DMBs to bivalent TSS/enhancer regions coincides with the ability of some pioneer TFs to simulate PRC2 complex-inducing H3K27me3-marked heterochromatin, often deposited on lineage-specific enhancers (**Supplemental Fig. 2d**) (66–68). In composite, these results match with the role of the liver cell-type-specific methylation blocks identified here as being critical for cell identity.

### Origins of cellular damage immediately after liver transplant

To identify the origins of cfDNA fragments in the circulation of liver transplant patients, we used the top 100 methylation blocks for each cell-type group and generated an expanded liver cell-type-specific DNA methylation atlas (Methods). We then applied a fragment-level deconvolution algorithm, previously validated to estimate relative contributions from cfDNA methylation sequencing data (**Supplemental Table 6a**) (35). First, to explore the changing cfDNA makeup after liver transplant, we assessed changes across all patients comparing pre-transplant cfDNA origins to post-reperfusion changes in serially collected blood samples from 28 liver transplant patients on the day of surgery (POD0; **Fig. 4a**). We found that there was a significant ∼5-fold increase in cfDNA concentration after transplant across all patients in the cohort, reflecting increased cell turnover from the surgical procedure itself (p<0.05, Wilcoxon matched-pairs signed rank test) (**Fig. 4f**). From the deconvolution analysis we found that liver cell types mainly contributed to this increase (**Fig. 4b**) with a significant increase in hepatocyte, hepatic stellate, and endothelial cfDNA fraction and a corresponding relative decrease in myeloid cfDNA that constitutes most of the hematopoietic signal at baseline (p<0.05, Wilcoxon matched-pairs signed rank test) (**Fig. 4d, e, g, h**). The concentration of hepatocyte cfDNA in genome equivalents/mL (Geq/mL) correlated with patient AST and ALT liver enzyme values (Spearman r = 0.81 AST and r = 0.82 ALT, p<0.05) (**Fig. 4c**). The homogenous cfDNA changes across all patients show applicability of this approach across a diverse patient cohort. Beyond damages to the allografted liver cell-types, we found that the transplant procedure results in multiple tissue cellular damages of the recipient as well. We were surprised to find a significant increase in neuron-derived cfDNA after transplant (**Fig. 4i**). While only representing a small overall fraction of the total cfDNA, there was an average 4-fold increase in this signal indicating neuronal cell death during the procedure. In addition, there were also significant increases in cardiomyocyte, biliary-ductal and gastric-epithelial cfDNAs in Geq/mL (**Supplemental Fig. 3**).

**Figure 4.**
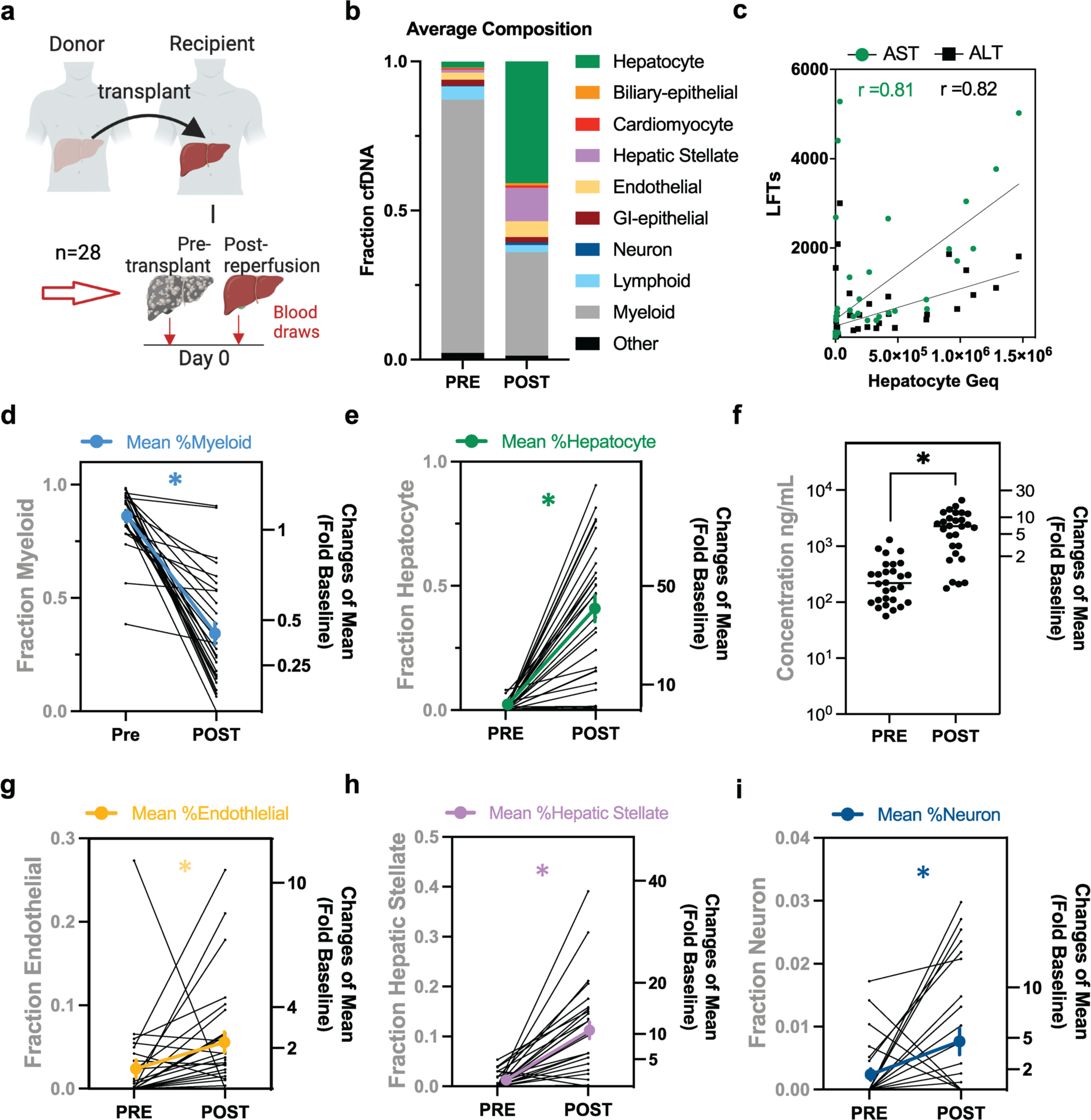
Origins of cellular damage after liver transplant derived from methylated cfDNA fragments. **a**, Serial serum samples from 28 liver transplant patients collected pre-transplant and post-reperfusion on post-operative day 0 (POD0). **b**, Average cellular origins of cfDNA estimates. **c**, Correlation of AST and ALT enzyme activity with hepatocyte-derived cfDNA (p<0.05, Spearman r = 0.81 AST; r = 0.82 ALT). **d**, **e**, **g**-**i**, Pre-transplant (PRE) and post-reperfusion (POST) fraction of cfDNAs from myeloid (d), hepatocyte (e), endothelial (g), hepatic stellate (h), and neuronal cells (i) of individual patients. The mean ± SEM of the cohort is shown in bold. Right axes: Fold change relative to pre-transplant. **f**, Concentration of cfDNA isolated from patient serum. Individual patient and median values are shown. Wilcoxon matched-pairs signed rank test was used for comparison amongst groups (d-i; n=28). *p<0.05; myeloid p=0.0001, hepatocyte p=0.0001, endothelial p=0.0025, hepatic stellate p=0.0001, neuron p=0.029, concentration p=0.001.

### Sustained elevation of hepatocyte and biliary epithelial cfDNA indicate allograft injury

We collected additional serum samples in a subset of 20 liver transplant patients to explore cfDNA changes over time during the first month after transplant, the highest risk period for post-transplant complications (**Fig. 5a**). Of these patients, 11 (55%) had liver biopsies showing allograft injury within the first year. There were no differences in cfDNA concentration after transplant comparing these two outcome groups (**Fig. 5B**) though we noticed changes in cfDNA composition when comparing across the entire cohort (**Supplemental Fig. 4a**). Liver-epithelial cellular damage mostly recovered in patients without allograft injury during the first post-operative week. In contrast, patients diagnosed with allograft injury during the first year after transplant had sustained elevation of hepatocyte and biliary epithelial cfDNA from POD7-POD30 (p<0.05, Mann-Whitney test) (**Fig. 5c to 5f**). Despite these differences in liver epithelial signals, there was no significant difference in hepatic stellate or endothelial cfDNA associated with different outcomes (**Supplemental Figs. 4b and 4c**). These findings were irrespective of the type of allograft injury diagnosed at the eventual time of for-cause liver biopsy (FC-bx). Of the patients with allograft injury, 7 (of 20) were diagnosed with hepatocellular, 3 mixed hepatobiliary, and 1 biliary forms of allograft injury. The majority of patients were diagnosed with allograft injury beyond the first month, but five patients were diagnosed within the first month (all within the first-year post-transplant). Despite variation in timing, elevated liver epithelial cfDNA was detected during the first post-operative month in all 11 patients with allograft injury, with an elevated signal detected a median of 63 days [range 2-203 days] ahead of the time of tissue-biopsy based diagnosis. Comparing the trajectory of liver cell-type damages over time, patients without allograft injury had higher levels of lymphoid and endothelial cfDNA relative to hepatocyte and biliary epithelial cfDNA at POD30, suggesting that the ratio may be useful to monitor and predict injury patterns during the peri-transplant time period (**Fig. 5g**).

**Figure 5.**
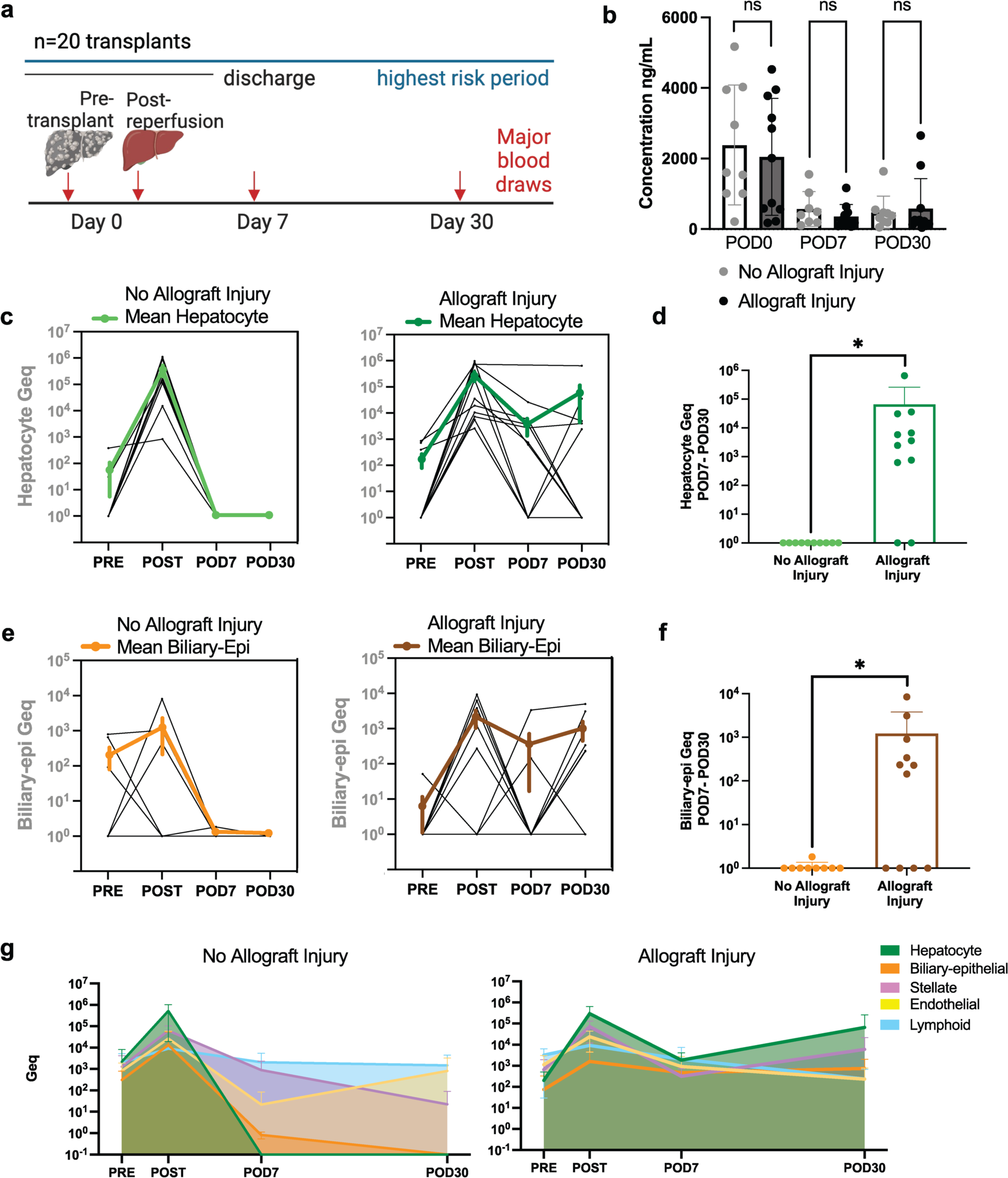
Time course of cell-type-specific damages after liver transplant. **a**, Serial serum samples from 20 liver transplant patients collected pre-transplant, post-reperfusion (POD0), post-operative day 7 and 30 (POD7, POD30). By 1 year after transplant, 11 of 20 patients were diagnosed with allograft injury by for-cause biopsy. **b**, Concentration of cfDNA isolated from patient serum. Individual values and mean ± SD at each timepoint, grouped by outcome. (Mann-Whitney test, ns p>0.05). **c**, Hepatocyte cfDNA time course in patients with allograft injury (right) or no allograft injury (left). Mean ± SEM of each cohort is shown in bold. **d**, Average hepatocyte cfDNA on POD7 and POD30, grouped by outcome (Mann-Whitney test, p<0.05). **e**, Biliary cfDNA time course in patients with allograft injury (right) or no allograft injury (left). Mean ± SEM of each cohort is shown in bold. **f**, Average biliary cfDNA on POD7 and POD30, grouped by outcome (Mann-Whitney test, p<0.05). **g**, Time course of five liver cell type cfDNAs in patients with allograft injury (right) or no allograft injury (left). Mean + SD. (**b,d,f**) NS p≥0.05, *p<0.05; concentration POD0 p=0.516; concentration POD7 p=0.237; concentration POD30 p=0.503; biliary p=0.009; hepatocyte p=0.002.

### Cell-free methylated DNA indicates the source of allograft injury

We added phenotype-matched samples from additional patients at the time of for-cause liver biopsy (FC-bx) used to diagnose allograft injury, evaluating 30 serum samples from 24 individuals (**Fig. 6a**). Samples were classified as having hepatocellular (n=14), biliary (n=6), or mixed hepatobiliary (n=10) forms of allograft injury from histopathological analysis of the paired biopsy tissues. Notably, the composition of cfDNA was significantly different at the time of biopsy-proven phenotypes, comparing hepatocellular and biliary etiologies of allograft injury (**Fig. 6b** and **Supplemental Table 6b**). Hepatocyte cfDNA was increased in samples with hepatocellular or mixed hepatobiliary injury compared to pure biliary injury (p<0.05, Mann-Whitney test) (**Fig. 6c**). Likewise, biliary cfDNA was increased in samples with biliary or mixed hepatobiliary injury compared to hepatocellular injury (p<0.05, Mann-Whitney test) (**Fig. 6d**). The cfDNA composition changes over time reflected the trajectory of cellular damages in patients with different injury types (**Fig. 6e to 6h**; patient details in the legend). At the time of FC-bx only hepatocyte cfDNA was detected in a patient with hepatocellular injury (**Fig. 6e**), only biliary epithelial cfDNA in a patient with pure biliary injury (**Fig. 6f**), and both hepatocyte and biliary epithelial cfDNA in two patients with different etiologies of mixed hepatobiliary injury (**Fig. 6g and 6h**). Taken as a whole, distinct cellular damages after liver transplant are detectable by the analysis of blood samples.

**Figure 6.**
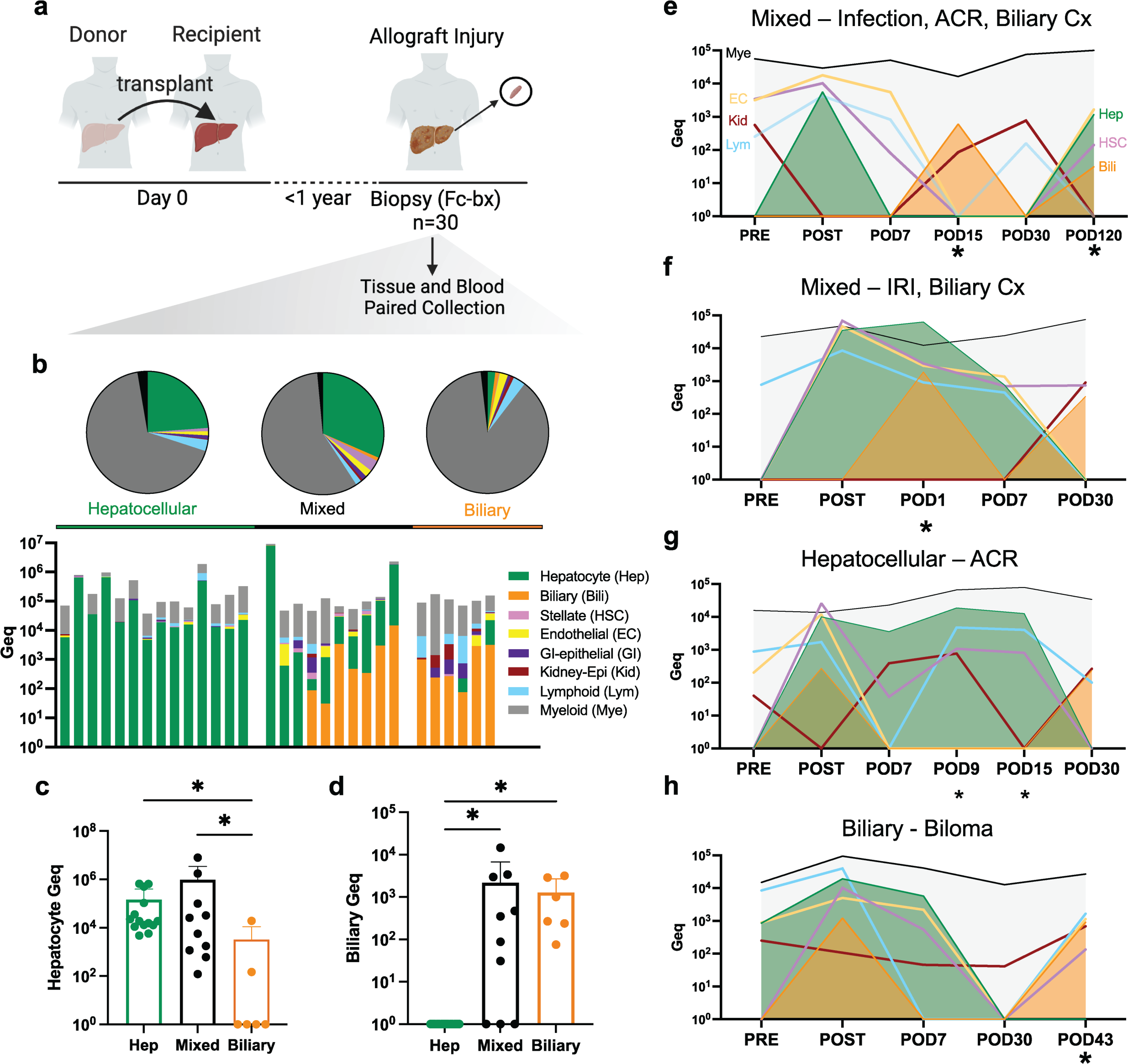
Cell-free methylated DNA indicates cellular sources of allograft injury. **a,** Serum samples collected at the time of for-cause liver biopsies (FC-bx) to diagnose allograft injury. All biopsies were taken within 1 year of liver transplant and samples are representative of 24 patients (n=30 samples). **b**, Cellular origins of cfDNAs classified by injury patterns observed in biopsies. Top, Average fractions of different cellular sources detected for each type of allograft injury. Bottom, Amount of cfDNAs (in Geqs) in individual patients. **c**, Hepatocyte cfDNA in serum samples with hepatocellular or mixed hepatobiliary injury compared to biliary injury alone (Mann-Whitney test, p<0.05). **d**, Biliary cfDNA in serum samples with biliary or mixed hepatobiliary injury compared to hepatocellular injury alone (Mann-Whitney test, p<0.05). **b**-**d**, Serum samples classified as n=14 hepatocellular, n=10 mixed hepatobiliary, and n=6 biliary etiologies of allograft injury. **e**-**h**, Time courses of cellular damage during the peri-transplant time period in patients with hepatocellular, biliary, and mixed hepatobiliary forms of allograft injury. Timepoints corresponding to complications and liver-biopsy proven diagnoses are marked by an asterix. **e**, Patient with COVID-19 infection at POD15 and FC-bx diagnosis of acute cellular rejection (ACR) with hyperbilirubinemia at POD120 (mixed injury classification). Elevated kidney epithelial cfDNA detected on POD0, POD15, and POD30 match with the hepato-renal syndrome (HRS) diagnosis pre-transplant and acute kidney injury (AKI) after transplant. **f**, Patient with FC-bx diagnosis of hepatic ischemia with hyperbilirubinemia at POD1 (mixed injury classification). AKI was indicated by elevated creatinine levels POD9 and elevated kidney epithelial cfDNA on POD30. **g**, Patient with FC-bx diagnosis of Acute Cellular Rejection (ACR) at POD9 and POD15 (hepatocellular injury classification). AKI was indicated by elevated creatinine levels on POD8 and elevated kidney epithelial cfDNA POD7, POD9, and POD30. **h**, Patient with diagnosis of biloma at POD43 (biliary injury classification).

## Discussion

The liver cell-type-specific DNA methylation atlas sheds light onto the epigenomic characteristics established early during development by identifying genomic regions of cell identity that are stably maintained in differentiated cells. We find that DNA methylation coincides with other liver cell-type-specific epigenetic marks, validating the biological relevance of these regions and enhancing their utility in detecting altered turnover of cells from DNA fragments shed in the circulation. Similar to other studies, we found the majority of liver-cell-specific DMBs to be hypomethylated. However, we found enriched TFBS of pioneer factors within both hypo- and hyper-methylated liver-cell-specific DMBs. Surprisingly, we also found CpG dinucleotides enriched within TFBS motifs associated with methylation-sensitive TFs in hypermethylated DMBs. This suggests that cell-type-specific hypo- and hyper-methylated regions may play a similar function in different contexts to repress precursor or stem cell transcriptional programs and control terminal differentiation into distinct cell types. Ultimately, this could serve as a valuable resource for many applications and shed light on factors needed for cell-reprogramming that may also play a role in disease pathogenesis and cell-type-specific epigenetic regulation (69).

We identified sufficient numbers of DNA methylation blocks to discriminate cells of origin of liver-derived DNA fragments in the circulation, including hepatocyte, biliary-epithelial, hepatic stellate, and endothelial cells. As one limitation, we were unable to identify enough DMBs with sufficient specificity to profile liver-resident immune cell turnover in patient blood samples relative to all other cell-types included in the atlas. Likely this was due to the purity and extent of peripheral and tissue-resident immune cell methylome reference data available. Instead, extended liver-resident immune cell markers were identified with relaxed specificity thresholds to use for characterization of liver cell-specific epigenetic data (**Supplemental Table 7** and Supplemental Methods). Generation of additional cell-specific methylation sequencing data to better characterize immune cell diversity will allow for enhanced ability to detect tissue-resident cell turnover in the future. In addition, implementation of hierarchical statistical models will allow for a more fine-tuned assessment of cell-free DNA composition in the face of limited numbers of highly specific methylation patterns to distinguish rare cell-types (70, 71).

The expanded liver cell-type-specific methylation atlas allowed for detection of tissue-derived fragments in the circulation to reveal cell types in the recipient impacted by the transplant procedure, comparing post-reperfusion signals to the pre-transplant baseline. The donor liver can be damaged in several ways including, cold and warm ischemia, surgical anastomoses, and reperfusion injuries (1–3, 33, 34). We found that these tissue effects were reflected by a relative increase in hepatocyte, hepatic stellate, and endothelial cfDNA compared to the myeloid-derived baseline signal. However, we were surprised by the increase in neuron and cardiomyocyte cfDNA after the transplant. The increase in neuron cfDNA could be caused by neurotoxicity from the general anesthesia; although, neurological complications are more common after liver (30%) than after heart (4%) or kidney transplants (0.5%) (72). Increased susceptibility or comorbidity due to the pathophysiology of the underlying hepatic disease may play a role and contribute to increased neuronal cell death in liver transplant patients.

The large increase in liver-cell-derived cfDNA after transplant can serve as a proof-of-concept to validate the prediction accuracy of deconvolution results. We used a complete fragment-level deconvolution model to estimate relative abundance changes to cfDNA composition, shown to accurately detect cfDNA from a source at 0.1% resolution (35). Also, we found significant correlation with hepatocyte cfDNA and AST/ALT liver enzyme activity (**Fig. 4c**). However, we did not find biliary cfDNA to correlate significantly with alkaline phosphatase (ALP) or bilirubin levels (**Supplemental Figs. 4d and 4e**). The short half-life of cfDNA (15 mins – 2 hours) relative to commonly monitored liver function parameters also contributes to some discrepancy between the observed values (73, 74). Changes in cell-type-specific cfDNAs reflect changes in cell turnover and thus measure different facets of tissue dysfunction.

We found that sustained elevation of liver epithelial cfDNA during the first month after transplant was associated with allograft injury, while patients without allograft injury had significantly reduced levels of liver epithelial cfDNA as early as the first week post-transplant. Importantly, liver cell-specific methylation patterns appear to be stably maintained during the ongoing processes of tissue damage, repair, and remodeling after transplant. We were able to detect elevated liver epithelial signals in all patients with allograft injury, despite being a diverse cohort with several different types of allograft injury represented. Our results suggest that cell-free methylated DNA has predictive and diagnostic value to detect allograft injury earlier than clinical diagnosis by liver biopsy. Notably, we did not find a significant difference in hepatic stellate or endothelial cfDNA comparing patients with allograft injury to those without allograft injury during the first month post-transplant. Interestingly, in liver damage after radiation treatment of patients with right-sided breast cancers liver endothelial cfDNA showed a >10-fold increase after radiation and delayed recovery to baseline one month after treatment in comparison to hepatocyte cfDNA (38). Thus, cfDNAs reflect distinct cellular responses to different types of injury and repair in the same organ.

Beyond damage to the allograft, methylated cfDNA is also able to reveal cellular damages of other recipient organs to indicate extra-hepatic toxicity and immune cell turnover (23, 30, 31). This is a useful application of methylated cfDNA that can simultaneously allow for monitoring of common pulmonary, renal, cardiac, and neurological complications after liver transplants. Acute kidney injury (AKI) is one of the most common post-operative complications, occurring in up to 78% of liver transplant patients (2). We were able to detect elevated kidney epithelial cfDNA in several patients in our cohort experiencing hepato-renal syndrome (HRS) pre-transplant as well as those experiencing AKI post-transplant (**Fig. 6e-6h**). In addition, we noticed divergent trajectories of lymphoid versus myeloid cfDNA, with lymphoid cfDNA demonstrating more dynamic changes compared to myeloid cfDNA that remains a constant background signal (**Fig. 6e-6h**). We found elevated lymphoid cfDNA corresponding to infection in several patients, including one patient with a COVID-19 infection (**Fig. 6e**).

Many studies have demonstrated the utility of donor-derived (dd) cfDNA to detect allograft injury (12, 14–17). However, dd-cfDNA is unable to discriminate amongst different causes of allograft injury. Likewise, it remains a challenge to distinguish causes relying on clinical presentation alone. Therefore, liver biopsy is still the gold standard to confirm a diagnosis and evaluate for response to treatment (4). Here we found that methylated cfDNA is able to detect and differentiate hepatocellular versus biliary causes of allograft injury at the time of biopsy-proven diagnosis (FC-bx). Biliary complications after liver transplant, such as ascending cholangitis, strictures (both anastomotic and non-anastomotic), leaks, and recurrence of primary sclerosing cholangitis, contribute significantly to post-transplant morbidity and mortality in both living and deceased donor transplant recipients. Conventional diagnostic and monitoring methods for these conditions often necessitate cross-sectional imaging techniques, such as MRCP, or invasive procedures like ERCP or liver biopsy, which pose additional risks to patients (75, 76). Enhanced detection of biliary cell-type-specific damage allows for differentiation from hepatocellular forms of allograft injury and associated tissue damage. This enables an earlier and more accurate diagnosis of biliary complications and improved non-invasive monitoring post-treatment. Incorporating cfDNA as a diagnostic tool into clinical practice could potentially reduce the need for invasive procedures and facilitate early intervention with targeted treatment.

In summary, we show that changes in methylated cfDNA released from dying cells can indicate increased cell death and tissue damage in transplant patients. Expanded atlases of DNA methylation sequencing data allow for identification of cfDNA fragments originating from a variety of cell-types in the liver, demonstrating applicability in a wide range of clinical settings. We correlate our findings from the methylated cfDNA analysis with clinical data, histopathological results, and outcomes of conventional clinical monitoring. We conclude that cell-free methylated DNA in the circulation of liver transplant patients can indicate allograft injury and discriminate amongst causes of allograft injury matching with tissue biopsy-proven diagnosis.

## Methods

### Study cohort

Serial serum samples were collected from 28 liver transplant patients at predetermined timepoints; pre-transplant (PRE) and post-reperfusion (POST) on post-operative day 0 (POD0), post-operative day 7 (POD0), and post-operative day 30 (POD30). Beyond this, we also collected samples in patients experiencing complications at the time of symptom presentation. Further, we added phenotype-matched samples from an additional 16 patients at the time of for-cause liver biopsy (FC-bx) used to diagnose allograft injury. Samples were classified as having hepatocellular (n=14), biliary (n=6), or mixed hepatobiliary (n=10) forms of allograft injury from histopathological analysis of paired liver biopsy tissues (additional details in Supplemental Methods section). A schematic of the time series for sample collection can be found in **Fig. 1**. For serum isolation, peripheral blood (∼6-12 mL) was collected in red-top venous puncture tubes and allowed to clot at room temperature for 30 minutes before centrifugation at 1200 x g for 10 min at room temperature to separate the serum fraction. Patient characteristics with samples analyzed in this study are summarized in **Supplemental Table 1.**

### Isolation of circulating cell-free DNA (cfDNA)

Circulating cell-free DNA was extracted from 2-6 mL human serum, using the QIAamp Circulating Nucleic Acid kit (Qiagen) according to the manufacturer’s instructions. Cell-free DNA was quantified via Qubit fluorometer using the dsDNA BR Assay Kit (Thermo Fisher Scientific). Additional size selection using Beckman Coulter beads was applied to remove high-molecular weight DNA reflective of cell-lysis and leukocyte contamination as previously described (38, 77). Paired serum and plasma were processed at serial timepoints for n=3 patients to serve as a quality control (**Supplemental Fig. 5** and Supplemental Methods). Fragment size distribution of isolated cfDNA after size selection was validated on the 2100 Bioanalyzer TapeStation (Agilent Technologies).

### Cell isolation to generate reference liver cell-type epigenomes

Reference epigenomes were generated for human liver cell types to expand upon publicly available datasets (**Supplemental Table 2**). Human biliary tissues were obtained from organs not suitable for transplant that were otherwise normal according to surgical assessment. Tissues were dissected, with samples processed from lobes, common hepatic duct, gallbladder, and common bile duct. Biliary epithelial cells (EpCAM+) were isolated from the dissected tissues according to previously established protocols (details in the Supplemental Methods section; **Supplemental Fig. 1b**) (78, 79). Cryopreserved passage 1 human liver sinusoidal endothelial cells (LSEC) were purchased from ScienCell research laboratories (SKU#5000). Cryopreserved passage 0 liver-resident immune cells (Kupffer) and passage 1 human hepatic stellate cells were isolated from single donor healthy human tissues purchased from Novabiosis Lot: QGJ and JNA (liver-immune); Lot: ZMC and WAP (hepatic-stellate). Paired RNA-seq data was generated from the same cell-populations used for DNA methylation profiling to validate the identity of cell-types obtained from commercial sources through analysis of cell type expression markers.

### Isolation and fragmentation of genomic DNA

Genomic DNA from tissues was extracted with the DNeasy Blood and Tissue Kit (Qiagen) following the manufacturer’s instructions and quantified via the Qubit fluorometer dsDNA BR Assay Kit (Thermo Fisher Scientific). Genomic DNA was fragmented via sonication using a Covaris M220 instrument to the recommended 150-200 base pairs before library preparation. Lambda phage DNA (Promega Corporation) was also fragmented and included as a spike-in to all DNA samples at 0.5%w/w, serving as an internal unmethylated control. Bisulfite conversion efficiency was calculated through assessing the number of unconverted C’s on unmethylated lambda phage DNA.

### Bisulfite capture-sequencing library preparation

Bisulfite capture-sequencing libraries were generated as previously described. In brief, WGBS libraries were generated using the Pico Methyl-Seq Library Prep Kit (Zymo Research) according to the manufacturer’s instructions. Library quality control was performed with an Agilent 2100 Bioanalyzer and quantity determined via the KAPA Library Quantification Kit (KAPA Biosystems). WGBS libraries were then pooled to meet the required 1µg DNA input necessary for targeted enrichment. However, no more than four WGBS libraries were pooled in a single hybridization reaction and the 1ug input DNA was divided evenly between the libraries to be multiplexed. Hybridization capture was carried out according to the SeqCap Epi Enrichment System protocol (Roche NimbleGen, Inc.) using SeqCap Epi CpGiant probe pools with xGen Universal Blocker-TS Mix (Integrated DNA Technologies, USA) as the blocking reagent. Washing and recovering of the captured library, as well as PCR amplification and final purification, were carried out as recommended by the manufacturer. The capture library products were assessed by Agilent Bioanalyzer DNA 1000 assays (Agilent Technologies, Inc.). Bisulfite capture-sequencing libraries with inclusion of 15-20% spike-in PhiX Control v3 library (Illumina) were clustered on an Illumina Novaseq 6000 S4 flow cell followed by 150bp paired-end sequencing.

### Bisulfite sequencing data alignment and preprocessing

Paired-end FASTQ files were trimmed using TrimGalore (V 0.6.6) (80) with parameters “--paired -q 20 --clip_R1 10 --clip_R2 10 --three_prime_clip_R1 10 --three_prime_clip_R2 10”. Trimmed paired-end FASTQ reads were mapped to the human genome (GRCh37/hg19 build) using Bismark (V 0.22.3) (81) with parameters “--non-directional”, then converted to BAM files using Samtools (V 1.12) (82). BAM files were sorted and indexed using Samtools (V1.12). Reads were stripped from non-CpG nucleotides and converted to BETA and PAT files using *wgbstools* (V 0.1.0) (https://github.com/nloyfer/wgbs_tools), a tool suite for working with WGBS data while preserving read-specific intrinsic dependencies (83). The BETA files (a wgbstools-compatible binary format) contain position and average methylation information for single CpG sites. The PAT files contain fragment-level information (including CpG starting index, methylation pattern of all covered CpGs and number of fragments with exact multiCpG pattern).

### Reference DNA methylation data from healthy tissues and cells

Availability of previously published and publicly available WGBS data from healthy cell-types and tissues used in this paper are described in **Supplemental Table 2**. Controlled access to reference WGBS data from normal human tissues and cell types were requested from public consortia participating in the International Human Epigenome Consortium (IHEC) (84) and upon approval downloaded from the European Genome-Phenome Archive (EGA), Japanese Genotype-phenotype Archive (JGA), database of Genotypes and Phenotypes (dbGAP), and ENCODE portal data repositories (85). Reference WGBS data were also downloaded from selected GEO and SRA datasets. Reference WGBS data were analyzed as previously described (38).

### Generation of expanded cell-type-specific DNA methylation atlas

Previously established atlases of cell-type-specific DNA methylation were refined to include expanded data generated from liver cell-types and curated from recently published WGBS dataset of purified healthy human cell-types (**Supplemental Table 2**). Tissue and cell-type specific methylation blocks were identified from reference WGBS as previously described (additional details in Supplemental Methods section) (38). In brief, data was first segmented into blocks of homogenous methylation and then analysis was restricted to blocks covered by the hybridization capture panel used in the analysis of cfDNA (probed regions span 80Mb (∼20% of CpGs) on the capture panel) (35). We also restricted analysis to blocks containing a minimum of three CpG sites, with lengths less than 2kb and at least 10 observations. Samples were divided into 20 groups by cell-type and we performed a one-vs-all comparison to identify differentially methylated blocks unique for each group. For this we used the find_markers Rscript (with parameters “--tg.quant 0.2 --bg.quant 0.1 --margin 0.4”) to calculate the average methylation per block/sample and rank the blocks according to the difference in average methylation between any sample from the target group and all other samples (38) (Supplemental Code). Blocks with a (-) direction are hypomethylated and (+) direction are hypermethylated, defined as a as a direction of methylation in the target cell-type relative to all other tissues and cell-types included in the atlas. Identified liver cell-type-specific DMBs meeting these specified requirements can be found in **Supplemental Table 3**. Extended cell-type-specific blocks for liver-resident immune cells can be found in **Supplemental Table 7** (additional details in Supplemental Methods section).

### UXM fragment-level deconvolution

An expanded atlas was generated using the top 100 methylation blocks per cell-type group using the ‘uxm build’ function with parameters “--rlen 3” from the UXM_deconv repository (https://github.com/nloyfer/UXM_deconv) (35). From this, each fragment is annotated as U (mostly unmethylated), M (mostly methylated) or X (mixed) depending on the number of methylated and unmethylated CpG sites. For each DMB, the proportion of U/X/M fragments is calculated across all reference WGBS cell-types and the U/M proportion reported for hypomethylated and hypermethylated DMBs, respectively. Then, the cell-type origins of cell-free DNA fragments isolated from serum of liver transplant patients are estimated using the ‘uxm deconv’ function with parameters “--rlen 3”. Briefly, a non-negative least squares (NNLS) algorithm is used to fit each cell-free DNA sample and estimate its relative contributions. Predicted cell-type proportions were converted to genome equivalents and reported as Geq/mL through multiplying the relative fraction of cell-type specific cfDNA times the concentration of cfDNA (ng/mL) by the mass of the human haploid genome 3.3 x 10^-12^ grams.

### Chromatin accessibility and histone modification data generation and analysis

ATAC-seq libraries were generated from liver sinusoidal endothelial (LSEC) and liver-resident immune cryopreserved cells using the ATAC-seq kit (Active Motif). H3K27ac histone modification data was generated from liver sinusoidal endothelial (LSEC) and liver-resident immune cryopreserved cells using the Cut&Tag-IT Assay kit (Active Motif, H3K27ac antibody cat#39133). Library products were assessed by Agilent Bioanalyzer HS DNA assays (Agilent Technologies, Inc.) and clustered on an Illumina Novaseq 6000 S4 flow cell followed by 150bp paired-end sequencing with inclusion of 10-15% spike-in PhiX Control v3 library (Illumina). Controlled access to DNase-seq, H3K4me1 and H3K27ac ChIP-seq data from hepatocytes was obtained from the German Epigenomme Program (DEEP) (EGAD00001002527) and publicly available ATAC-seq data from biliary tissue (gallbladder) was downloaded from the ENCODE project (ENCSR695FLC). Imputed data on H3K27ac binding in hepatic stellate cells was downloaded from the ENCODE project in bigWig format (ENCSR225UAF). ChromHMM annotations (18-state version) were downloaded from the ENCODE project (hepatocyte: ENCSR075JST, hepatic stellate: ENCSR593ZNP, and endothelial: ENCSR227ZSK) and genomic regions associated with H3K4me1 enhancer mark were extracted and reformatted in bigWig format. Paired-end FASTQ files were trimmed using TrimGalore (V 0.6.6) with parameters “--paired -q 20”. Trimmed paired-end FASTQ reads were mapped to the human genome (GRCh37/hg19 build) using Bowtie2-align (V 2.3.5.1) (86) with parameters “-X 1000 -- local --very-sensitive --no-mixed --dovetail --phred33”. Duplicated reads were marked with Picard (V 2.18.14) and reads with low mapping quality, duplicated or not mapped in proper pair were excluded using Samtools view (V 1.12) (82) with parameters “-F 1796 - q 30”. BAM files were sorted and indexed using Samtools. Chromatin accessibility data was normalized to the same depth and then bigWig files were created using deepTools (V 3.5.1) (87) with the functions ‘multiBamSummary ‘ and ‘bamCoverage’ using parameters “--normalizeUsing RPKM --binSize 25”. Detection q-values were calculated for histone modification data relative to Input control using the macs3 (V3.0.0a6) (88) bdgcmp function (-m qpois). This method uses the BH process for poisson p-values to calculate the score in any bin using control(Input) as lambda and treatment (IP sample) as observation. BigWig files were then generated using bedGraphToBigWig. Summary plots were prepared using deepTools (V 3.5.1) with functions ‘computeMatrix’ and ‘plotProfile’ using default parameters, except for ‘referencePoint=center’ and 5Kb margins.

### Functional annotation, transcription factor binding site and pathway analysis

Cell-type specific methylation blocks were annotated and transcription factor binding site analysis was performed using HOMER (V4.11.1) (89) and the annotatePeaks.pl and findMotifsGenome.pl functions (details in the Supplemental Methods section). Pathway analysis of genes adjacent to identified tissue and cell-type specific methylation blocks was performed using Ingenuity Pathway Analysis (IPA) (Qiagen) (90).

### Genome Browser and fragment-level visualizations

Reference WGBS samples were uploaded as custom tracks for visualization on the UCSC genome browser (91). Methylomes were converted to bigWig format using the wgbstools beta2bw function. Fragment-level visualization of methylation sequencing reads was performed with the wgbstools vis function with parameters “—min 3 –yebl -- strict”. Chromatin accessibility, histone modification, and RNA expression data were downloaded from the IHEC data portal as bigWig files (hg19). FOXA1 ChIP-seq data in liver was downloaded from the ENCODE project (ENCSR735KEY). Samples of the same cell type were averaged using multiBigwigSummary (v.3.5.1).

### Statistics

Statistical analyses for group comparisons and correlations were performed using Prism (GraphPad Software, Inc., United States) and R (V 4.1.3). A correlation analysis was performed to assess relationship between changing cell-free methylated DNAs and LFTs using Spearman’s Rank Correlation Coefficient. Statistically significant comparisons are shown, with significance defined as p<0.05. Correction for multiple hypothesis comparisons was performed using the Benjamini-Hochberg (B-H) method corrected p-value to control the false discovery rate (FDR) from multiple pathways being tested against each gene-set. A two-stage linear step-up procedure of Benjamini, Krieger and Yekutieli was performed for p-value adjustment from multiple outcome measures.

### Study Approval

Liver transplant patients were enrolled and provided signed informed consent in this Georgetown University Medical Center and MedStar Georgetown University Hospital IRB-approved study (IRB protocols # IRB 2017-0690 and # 2017-0365).

## Declarations

### Data availability

All data and code will be made available upon publication of this manuscript.

### Author Contributions

KO, VM, YC, DP and AHKK facilitated collection and processing of enrolled liver transplant patient samples and corresponding clinical data. MEM performed bisulfite-capture sequencing library preparation. MEM and AJK purified cell-types and generated reference liver cell-type-specific epigenome data. AJK and MOS performed RNA-sequencing analysis. MEM and SSJ performed bioinformatics analysis of NGS data. APM advised on statistical and computational data analysis. MEM and AW wrote the manuscript and generated all figures. MEM, AW, AR and AHKK conceived the study design and provided interpretation of results. All authors critically reviewed and approved the manuscript.

### Inclusion and Ethics statement

All collaborators of this study have fulfilled the criteria for authorship required by Nature Portfolio journals and have been included as authors, as their participation was essential for the design and implementation of the study. Roles and responsibilities were agreed among collaborators ahead of the research. This work includes findings that are locally relevant, which have been determined in collaboration with local partners. This research was not severely restricted or prohibited in the setting of the researchers, and does not result in stigmatization, incrimination, discrimination or personal risk to participants.

## Acknowledgements

This works was supported in part by funding from the National Institutes of Health USA (T32-CA009686 to MEM, F30-CA250307 to MEM, R01-AI132389 to AHKK, R01-CA231291 to AW and P30-CA51008 to AW). The graphical abstract and sample collection schemas in this manuscript were created with BioRender.com.

We also acknowledge the following consortia that generated data used in this study: *KNIH -* This study makes use of data generated by the Korea Epigenome Project. A full list of the investigators who contributed to the generation of the data is available from 152.99.75.168/KEP. Funding for the project was provided by KOREA EPIGENOME PROJECT.

*AMED-CREST / NBDC -* A part of the data used for this research was originally obtained by AMED-CREST International Human Epigenome Consortium (IHEC) research project/group led by Prof. /Dr. Yae Kanai and by Core Research and Evolutional Science and Technology (CREST), International Human Epigenome Consortium (IHEC) project/group led by Hiroyuki Sasaki, both available at the website of the National Bioscience Database Center (NBDC; http://biosciencedbc.jp/en/) of the Japan Science and Technology Agency (JST).

*CUHK CNARG -* This study makes use of data generated by The Chinese University of Hong Kong (CUHK) Circulating Nucleic Acids Research Group, as reported by Cheng THT et al. Clin Chem. 2019 Jul;65(7):927-936).

*ENCODE -* We downloaded the call sets form the ENCODE portal (https://www.encodeproject.org/) with the following identifiers: ENCSR656TQD, ENCSR328TBS, ENCSR116JEF, ENCSR116JEF, ENCSR999RWT, ENCSR577RUU, ENCSR267SNS, ENCSR899RXH, ENCSR258MDR, ENCSR641SDF, ENCSR835OJU, ENCSR442FIY, ENCSR893RHD, ENCSR784VGW, ENCSR211VXF, ENCSR749COU, ENCSR733ZTZ, ENCSR728AGC.

*CEEHRC -* The results published here are in part based upon data generated by The Canadian Epigenetics, Epigenomics, Environment and Health Research Consortium (CEEHRC) initiative funded by the Canadian Institutes of Health Research (CIHR), Genome BC, and Genome Quebec. Information about CEEHRC and the participating investigators and institutions can be found at http://www.cihr-irsc.gc.ca/e/43734.html.

*DEEP -* This study makes use of data generated by the Deutsches Epigenom Programm DEEP. A full list of the investigators who contributed to the generation of the data is available from the consortium website (www.deutsches-epigenom-programm.de).

*Blueprint -* This study makes use of data generated by the Blueprint Consortium. A full list of the investigators who contributed to the generation of the data is available from www.blueprint-epigenome.eu. Funding for the project was provided by the European Union’s Seventh Framework Programme (FP7/2007-2013) under grant agreement no 282510 – BLUEPRINT.

