## Supplemental Information for "Circulating, cell-free methylated DNA indicates cellular sources of allograft injury after liver transplant"

**Contents:**

Supplemental Materials and Methods

Supplemental References

Supplemental Figures and Legends

Legends for Supplemental Tables

#### Supplemental Materials and Methods

*Isolation of plasma samples*

Peripheral blood (~6-9 mL) was collected in lavender-top venous puncture tubes (EDTA tubes). Plasma (~8-12 mL) was collected as a part of the Ficoll-Paque density gradient separation method to isolate Peripheral Blood Mononuclear Cells (PBMCs) centrifuged at 1000 x g for 20 min at room temperature without brakes. The separate plasma fraction was then centrifuged again at 3000 x g for 10 min before cell-free DNA (cfDNA) isolation. CfDNA was isolated from plasma using the QIAamp Circulating Nucleic Acid kit (Qiagen) according to the manufacturer's instructions. Cell-free DNA was quantified via Qubit fluorometer using the dsDNA BR Assay Kit (Thermo Fisher Scientific).

*Comparison of methylation status in cfDNA isolated from paired serum and plasma samples*

Paired plasma and serum samples were collected from three liver transplant patients at serial timepoints to compare results across sample preparations. We computed the average methylation for each block and sample using wgbstools (--beta_to_table) (1). Correlation analysis was performed comparing the methylation status at the block level between the paired plasma and serum samples (**Supplemental Fig. 5**). Deconvolution analysis was performed to compare predicted cell-type proportions.

*Dissociation protocol for biliary epithelial tissues*

Human biliary tissues were obtained from organs not suitable for transplant that were otherwise normal according to surgical assessment. Tissues were dissected, with samples processed from lobes, common hepatic duct, gallbladder, and common bile duct. Biliary epithelial cells (EpCAM+) were isolated from the dissected tissues according to previously established protocols with some modifications (2, 3). The inner epithelial layer was separated from the outer fibrous connective tissue and muscle layer using disposable sterile scalpels and dissecting scissors. The inner epithelial layer was then minced and enzymatically digested in a solution of Collagenase (2 mg/mL, Roche)- DNase (0.5%,ThermoFisher)- Fetal Bovine Serum (2% FBS, GIBCO) in Advanced DMEM/F-12 (Invitrogen) supplemented with 1% Penicillin/Streptomycin (Pen/Strep, GIBCO) and 1% L-Glutamine (GIBCO). Using a set 1 hour program (37C_h_TDK3) on the GentleMACS Octo Dissociator (Miltenyi Biotec), tissues were further digested by transferring solution into gentleMACS C-tubes (Miltenyi Biotec). After digestion, samples were passed through a 70 μm cell strainer (Corning) and washed with DMEM/F-12 (Invitrogen). Samples were centrifuged at 300xg for 10 mins and the supernatant removed. Cells from intrahepatic ducts (lobes) were underlayed with an equal volume of 20% and 50% (v/v) Percoll (Sigma). Following centrifugation at 1800xg for 30 min at 4°C, the intrahepatic biliary epithelial (IHBEC) fraction at the interface of the 20% and 50% Percoll layers was collected, washed and re-suspended in EasySep bead buffer (PBS, 0.5% BSA, 0.5M EDTA). Biliary epithelial cells (EpCAM+) were isolated from pelleted cells using EasySep magnetic beads for EpCAM positive selection (StemCell Technologies Cat #17846) according to the manufacturer's protocol. Retained cells were considered EpCAM+

biliary epithelial cells. Efficiency of isolation was determined using flow cytometry (FITC Anti-human Epithelial cell, clone 5E11.3.1; StemCell Technologies Cat #60147FI).

*Classification of serum samples at time of for-cause liver biopsy (FC-bx) to diagnose graft injury*

Serum samples from 24 liver transplant patients (n = 30 serum samples) were taken at the time of for-cause liver biopsy (FC-bx) to diagnose graft injury. Serum samples were classified as having hepatocellular (n=14), biliary (n=6), or mixed hepatobiliary (n=10) forms of graft injury from histopathological analysis of paired liver biopsy tissues annotated by a pathologist. The following were defined as clinical etiologies of hepatocellular injury: Acute cellular rejection (ACR, rejection activity index, RAI 3+), recurrence of primary hepatic disease (including recurrence of viral hepatitis (HBV or HCV), autoimmune hepatitis, or (non)-alcoholic steatohepatitis in the transplanted organ), drug-induced hepatotoxicity, ischemia-reperfusion injury (IRI) leading to ischemic hepatitis. The following were defined as clinical etiologies of biliary injury: Anastomotic and non-anastomotic biliary strictures, ascending cholangitis, ischemic cholangiopathy, recurrence of primary biliary disease (including primary sclerosing cholangitis (PSC) among others), and septic cholestasis (4–8). Mixed hepatobiliary forms of graft injury were characterized by diagnosis of one or more hepatocellular and one or more biliary forms of graft injury at the same timepoint.

*Reference DNA methylation data from healthy tissues and cells*

Availability of previously published and publicly available WGBS data from healthy cell-types and tissues used in this paper are described in **Supplemental Table 2**. Controlled access to reference WGBS data from normal human tissues and cell types were requested from public consortia participating in the International Human Epigenome Consortium (IHEC) (9) and upon approval downloaded from the European Genome-Phenome Archive (EGA), Japanese Genotype-phenotype Archive (JGA), database of Genotypes and Phenotypes (dbGAP), and ENCODE portal data repositories (10–14). Reference WGBS data were also downloaded from selected GEO and SRA datasets (15–22). Reference WGBS data were analyzed as previously described.

*Segmentation and clustering analysis*

We segmented the genome into blocks of homogenous methylation as previously described (21). In brief, a Dynamic Programming segmentation algorithm was used to divide the genome into continuous genomic regions (blocks) showing homogenous methylation levels across multiple CpGs for each sample. We applied the segmentation algorithm to over 450 human reference WGBS methylomes and retained 364,268 blocks covered by the hybridization capture panel used in the analysis of cfDNA (probed regions span 80Mb (~20% of CpGs) on the capture panel). The top 10% most variable methylation blocks containing at least three CpG sites and coverage across 90% of samples were selected, irrespective of sample cell-type group. We computed the average methylation for each block and sample using wgbstools (--beta_to_table). Dimensional reduction was performed on the selected blocks using the UMAP package (V 0.2.8.2.0)

(23). Default UMAP parameters were used (15 neighbors, 2 components, Euclidean metric, and a minimum distance of 0.1).

*Identification of cell-type specific methylation blocks*

Tissue and cell-type specific methylation blocks were identified from reference WGBS as previously described (22). We performed a one-vs-all comparison to identify differentially methylated blocks unique for each group. All cell-type-specific blocks contained a minimum of three CpG sites, with lengths of less than 2kb and at least 10 observations. In brief, we calculated the average methylation per block/sample, as the ratio of methylated CpG observations across all sequenced reads from that block. Differential blocks were sorted by the margin of separation, termed "delta beta", defined as the minimal difference between the average methylation in any sample from the target group vs all other samples. Then, we computed the "soft margin" between target samples and background samples, allowing for some outliers using percentiles. For all hypomethylation markers we calculated the difference between the 80$^{th}$ percentile of the methylation status in the target group (--target.quant 0.2) and the 10$^{th}$ percentile of the methylation status in the background group (--bg.quant 0.1). Conversely, for all hypermethylated markers we calculated the difference between the 20$^{th}$ percentile methylation status in the target group and the 90$^{th}$ percentile in the background group. We selected blocks with a margin ≥ 0.4 for all cell-type groups. Blocks with a (-) direction are hypomethylated and (+) direction are hypermethylated, defined as a as a direction of methylation in the target cell-type relative to all other tissues and cell-types included in the atlas. We also used a magnitude threshold where all hypomethylated blocks have an

Average Methylation Fraction (AMF) <0.5 and hypermethylated blocks have an AMF >0.5. However, the vast majority of cell-type specific differentially methylated blocks are much more diverged (mean AMF hypo = <10% methylation and mean AMF hyper = >80%).

Biliary epithelial samples were separated into two groups of epithelial populations, the columnar intrahepatic and gallbladder versus cuboidal larger duct epithelial layers. DMBs were identified for each biliary epithelial cell population. Estimated cell proportions from columnar and cuboidal biliary epithelial cell proportions were combined to reflect the total biliary cfDNA from deconvolution analysis of serum samples from liver transplant patients. Endothelial samples were combined to identify common endothelial DMBs across all tissues. However, we ensured that methylation status was conserved in liver sinusoidal endothelial methylomes for all identified common endothelial-specific methylation blocks. Liver-resident immune (Kupffer CD14+,CD11b+,CD68+) samples were grouped with other myeloid immune samples when identifying markers for deconvolution of cfDNA fragments from liver transplant samples. Only two liver-resident immune cell-specific DMBs were identified when these samples were considered as a separate group at the same thresholds that other cell-type-specific DMBs were identified in the atlas. However, extended liver-resident immune cell markers were identified with relaxed thresholds (--margin 0.3, --target.quant 0.2, --bg.quant 0.2) to use for characterization of liver cell-specific epigenetic data. Chromatin accessibility and histone modification data in Figure 3A-C is plotted using identified biliary columnar epithelial, common endothelial, and extended liver-resident immune cell-specific DMBs (**Supplemental Tables 3 and 7**).

*Methylation score and visualization of cell-type specific methylation atlas*

Each DNA fragment was characterized as U (mostly unmethylated), M (mostly methylated) or X (mixed) based on the fraction of methylated CpG sites as previously described (21). We used thresholds of ≤33% methylated CpGs for U reads and ≥66% methylated CpGs for M. We then calculated a methylation score for each identified celltype specific block based on the proportion of U/X/M reads among all reads. The U proportion was used to define hypomethylated blocks and the M proportion was used to define hypermethylated blocks. Heatmaps were generated using the pretty heatmap function in the RStudio Package for the R bioconductor (RStudioTeam, 2015).

*Transcription factor binding site analysis*

Transcription factor binding site analysis was performed using the HOMER (V4.11.1) (24) findMotifsGenome.pl function for known and de novo motifs with parameters "-mask -chopify -size given -cpg" and with captured blocks without liver cell-type-specific methylation used as background. All DMBs for individual liver cell-type-specific hypomethylated DMBs were assessed. Similar analysis was performed comparing all hypomethylated DMBs and all hypermethylated DMBs for liver cell-types combined. Individual liver cell-type-specific hypermethylated DMBs were only assessed for hepatocytes, hepatic stellate, and biliary epithelial cell-types (due to limited hypermethylated DMBs identified for endothelial cells). For this analysis comparing all hyper- to all hypo- DMBs for liver cell types combined, enriched motifs containing CpG dinucleotides were quantified. All motifs with binomial p-value <0.05 were considered.

*RNA isolation and RNA-sequencing analysis*

RNA was isolated from sorted cells using the RNeasy Kit (Qiagen) according to the manufacturer's protocol and quantified by Qubit RNA BR assay (Thermo Fisher Scientific). Total RNA was validated using an Agilent RNA 6000 nano assay on the 2100 Bioanalyzer TapeStation (Agilent Technologies). The resulting RNA Integrity number (RIN) of samples selected for RNAseq analysis was at least 7. RNA-sequencing libraries were prepared using TruSeq Total RNA library Prep Kit (Illumina) at Novogene Corporation Inc., and 150bp paired-end sequencing was performed on an Illumina HiSeq 4000 with a depth of 50 million reads per sample. A reference index was generated using GTF annotation from GENCODEv28. Raw FASTQ files were aligned and assembled to GRCh38 and GRCh37 with HISAT2 / Stringtie (V 2.1.0) (25). The differential expression was analyzed in R with packages EdgeR (V 3.32.1) and Rsubread (V1.6.3) (26–28). Derived counts per million and p-values were used to create a rank ordered list, which was then used for subsequent integrative analysis. Expression levels at known cell type markers from single cell expression databases were used to validate the identity of isolated cell type populations for methylome analysis.

### Supplemental Figure 1

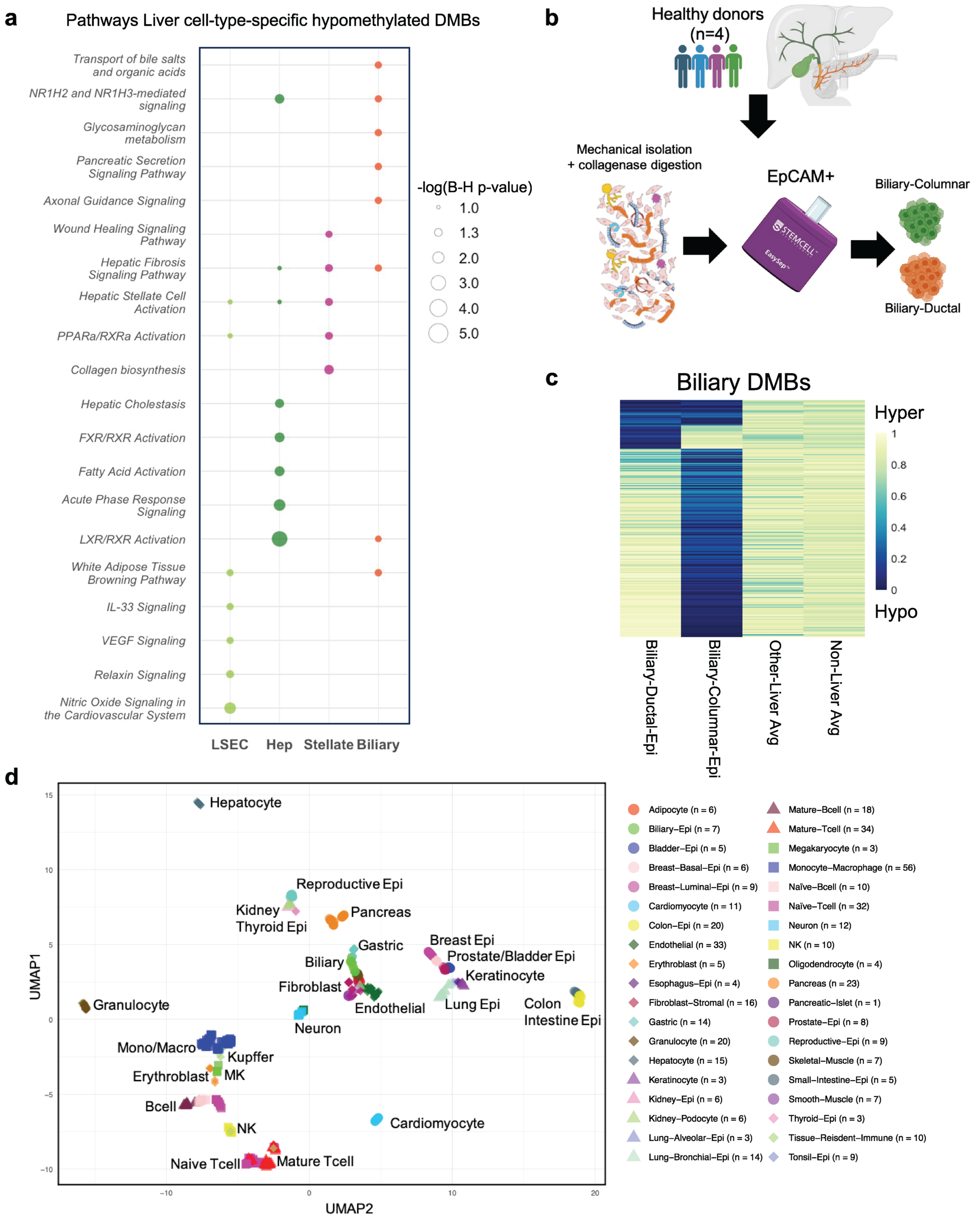

#### Supplemental Figures and Legends

**Supplemental Figure 1. Characterization of human healthy cell-type-specific reference methylation data. a**, Significant pathways related to the biological function of genes annotated to liver cell-type-specific hypomethylated blocks. **b**, Digestion of biliary tissues and enrichment of EpCAM (+) epithelial populations from the columnar epithelial versus ductal (cuboidal) epithelial layers. **c**, Heatmap of biliary epithelial DMBs. Each cell shows the average methylation across all CpGs in the block comparing biliary ductal (cuboidal epithelium), biliary columnar epithelial, average across other liver cell-types, and average across all non-liver cell-types in the atlas. **d**, UMAP projection depicting relationship between different cell types with WGBS reference datasets included for analysis. Average methylation was calculated for each sample within blocks of at least three CpG sites and the top 10% of captured blocks were selected showing the highest variability across all samples.

### Supplemental Figure 2

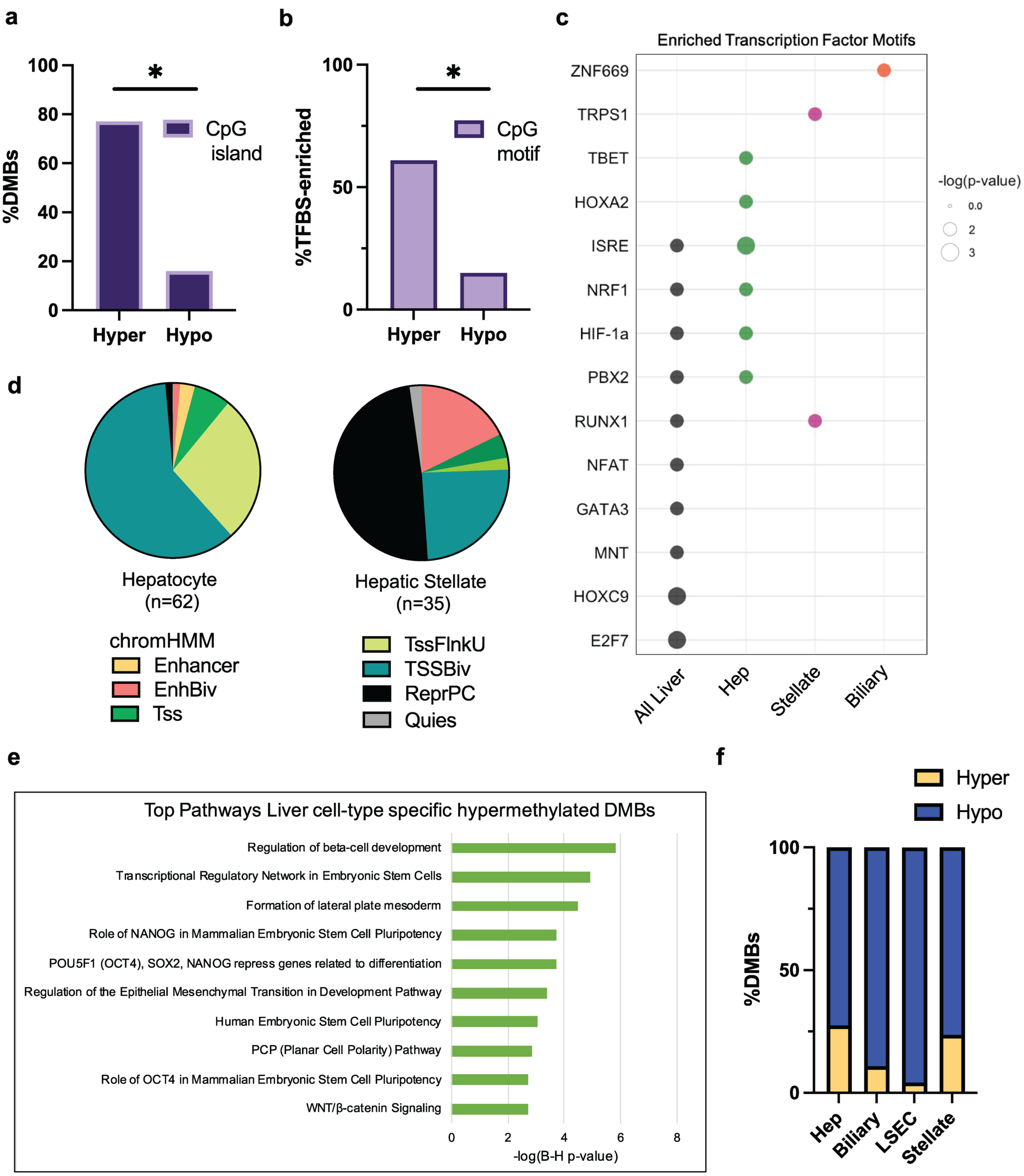

**Supplemental Figure2. Characterization of liver cell-type-specific hypermethylated DNA blocks.** **a**, Percent of liver cell-type-specific blocks that overlap with CpG islands based on UCSC hg19 annotations (77% hypermethylated; 16% hypomethylated blocks) (Fisher's exact test, $p<0.05$). **b**, Percent of motifs ($p <0.05$) enriched in liver cell-type-specific blocks containing a CpG dinucleotide (61% hypermethylated; 15% hypomethylated) (Fisher's exact test, $p<0.05$). **c**, Top TF binding sites enriched within liver cell type-specific hypermethylated blocks, from HOMER known motif analysis. Captured blocks without liver cell type-specific methylation were used as background. **d,** Fraction of cell type-specific hypermethylated blocks labeled as different chromatin states in chromHMM annotations from the same cell-type. **e**, Top 10 pathways related to the biological function of genes annotated to liver cell-type-specific hypermethylated blocks. **f**, Percent of hyper- and hypo-methylated DMBs identified for each liver cell type.

### Supplemental Figure 3

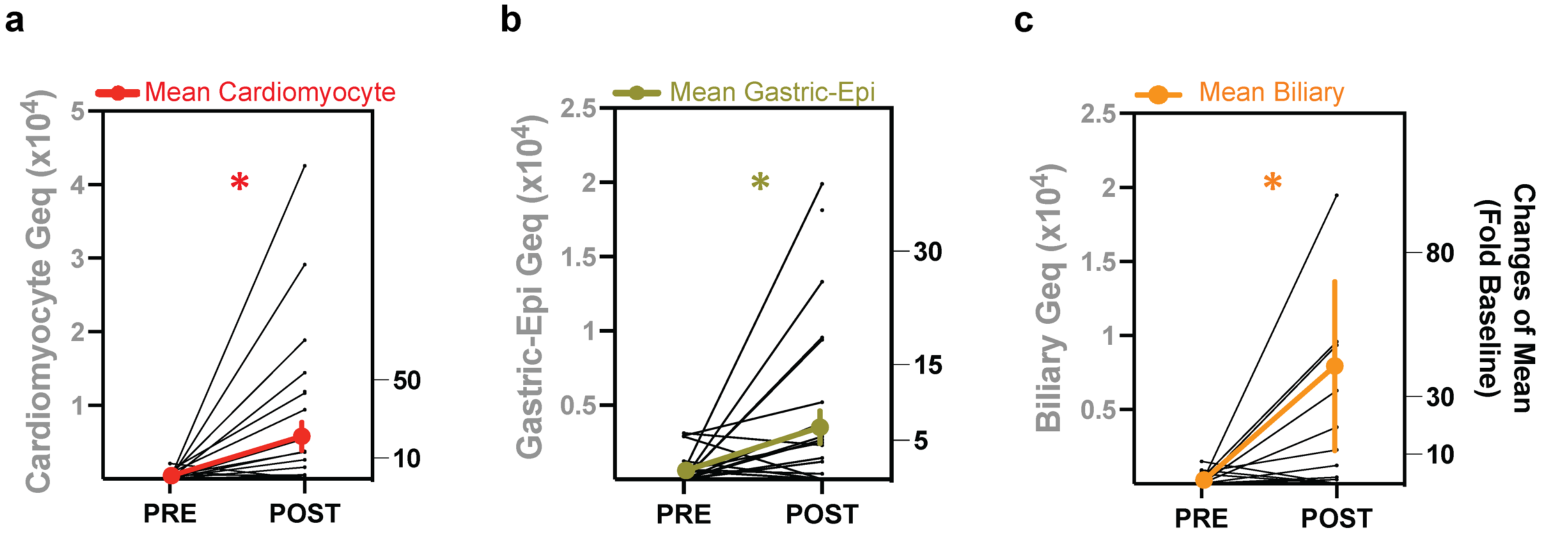

**Supplemental Figure 3. Expanded liver cell-type-specific DNA methylation atlases inform origins of cellular damage after liver transplant. a-c**, Cardiomyocyte, gastric-epithelial, and biliary-epithelial cfDNA (in Geq/mL) in serum samples collected pre-transplant (PRE) or post-reperfusion (POST) on POD0 (n=28 patients). Mean ± SEM fold change relative to pre-transplant levels is shown in bold. Wilcoxon matched-pairs signed rank test was used for comparison amongst groups. *$p<0.05$; cardiomyocyte $p=0.0027$, gastric-epithelial $p=0.015$, and biliary $p=0.0250$.

### Supplemental Figure 4

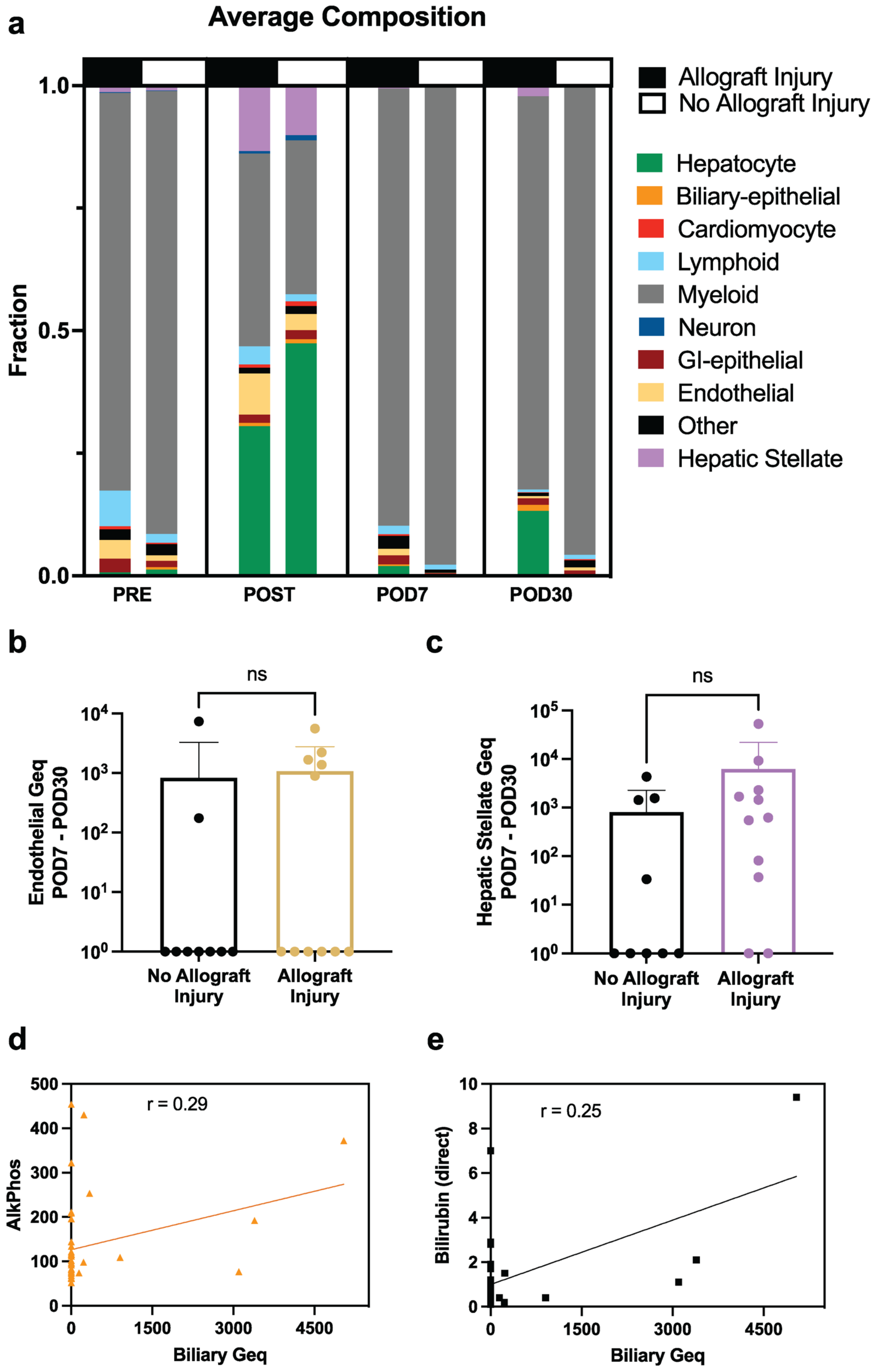

**Supplemental Figure 4. Cell-free DNA composition changes after transplant in patients with graft acceptance or injury. a**, Average cfDNA composition estimated from fragment-level deconvolution of serial serum samples collected from liver transplant patients pre-transplant (PRE) and post-reperfusion (POST) on day of transplant (POD0), post-operative day 7 (POD7), and post-operative day 30 (POD30). **b**, Average of endothelial cfDNA on POD7 and POD30 (Mann-Whitney test, ns $p>0.05$). **c**, Average of hepatic stellate cfDNA on POD7 and POD30 (Mann-Whitney test, ns $p>0.05$). **a-c**, Serum samples from 20 liver transplant patients collected pre-transplant, post-reperfusion (POD0), post-operative day 7 and 30 (POD7, POD30). By 6 months post-transplant 9 patients showed graft acceptance and 11 patients graft injury. **d**, **e**, Correlation of biliary cfDNA (biliary columnar epithelial + biliary cuboidal (ductal) epithelial) with alkaline phosphatase (ALP) serum levels (d; Spearman $r = 0.29$; ns $p>0.05$) or with serum bilirubin levels (e; Spearman $r = 0.25$; ns $p>0.05$).

### Supplemental Figure 5

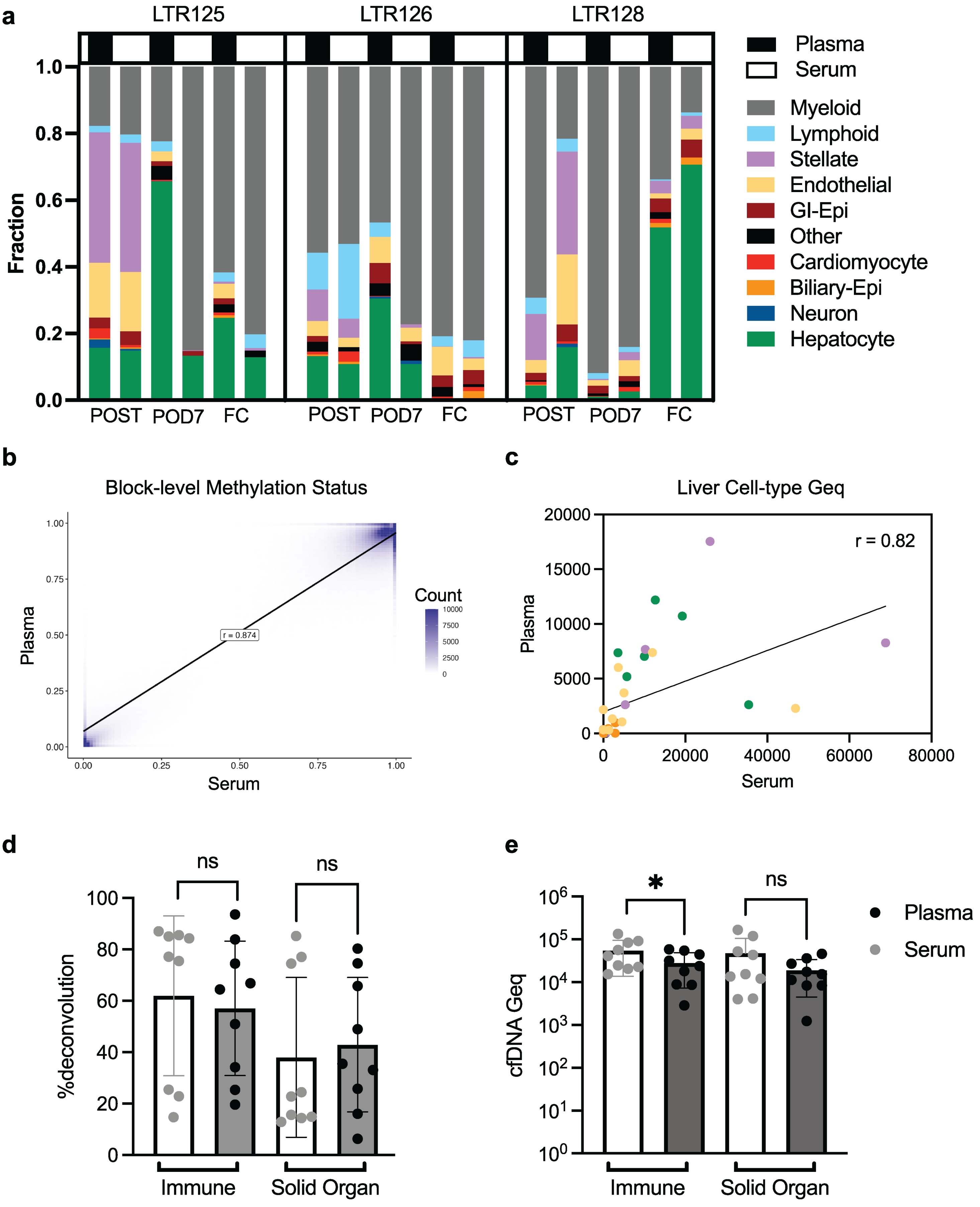

**Supplemental Figure 5. Comparison of methylation status and cellular origins of cfDNA isolated from serum and plasma of liver transplant patients. a.** Cellular origins of cfDNA fragments in paired serum and plasma samples (n=3 patients). **b.** Density heatmap comparing methylation status across blocks in cfDNA isolated from paired human serum and plasma (n=9 paired samples). Methylation status is represented by M-values (Logit transformation of $\beta$-values) that have normal distribution. Methylation levels are highly correlated at the block level with Pearson's r =0.874, $p < 0.05$). **c.** Correlation of predicted liver cell-type Geq in paired serum and plasma samples (spearman r = 0.82, $p < 0.05$). **d.** Predicted %Immune versus %Solid Organ derived cfDNA extracted from either serum or plasma. **e.** Immune and solid organ Geq from cfDNA isolated from serum versus plasma. **(d,e)** Data presented as mean $\pm$ SD; n=9 samples per group. Wilcoxon matched-pairs signed rank test was used for comparisons amongst groups. NS, $p \geq 0.05$; *$p < 0.05$.

#### Legends for Supplemental Tables

**Supplemental Table 1.** Characteristics of liver transplant patients enrolled in this study.

**Supplemental Table 2.** Human reference methylation data from healthy tissues and cell types.

**Supplemental Table 3.** Identified human liver cell-type-specific methylation blocks (margin 0.4;bg.quant 0.1;tg.quant 0.2). Annotation was performed using Homer. The margin of separation represents the delta-beta (maximum higher – minimum lower) across all samples. Blocks with a (-) direction are hypomethylated and (+) direction are hypermethylated. AMF (average methylation fraction) indicated as a fraction.

**Supplemental Table 4.** Significantly enriched biological pathways for genes associated with differential methylation in each cell-type.

**Supplemental Table 5.** Significantly enriched motifs from transcription factor binding site analysis (using HOMER). (a) Enriched motifs for individual liver cell-type-specific hypomethylated blocks (b) Enriched motifs for all hypomethylated DMBs in all liver cell-types combined. (c) Enriched motifs for all hypermethylated DMBs in all liver cell-types combined.

**Supplemental Table 6.** (a) Liver transplant serial cfDNA sample concentrations and predicted cell-type proportions from fragment-level deconvolution analysis (n=28 patients; n=100 samples). (b) Liver transplant phenotype-matched cfDNA sample concentrations and predicted cell-type proportions at FC-bx from fragment-level deconvolution analysis (additional n=16 patients; n=30 samples).

**Supplemental Table 7.** Extended liver-resident immune cell-type-specific methylation blocks (margin 0.3;bg.quant 0.2;tg.quant 0.2). Annotation was performed using Homer. The margin of separation represents the delta-beta (maximum higher – minimum lower) across all samples. Blocks with a (-) direction are hypomethylated and (+) direction are hypermethylated. AMF (average methylation fraction) indicated as a fraction.
